# Microscopic deconstruction of cortical circuit stimulation by transcranial ultrasound

**DOI:** 10.1101/2024.10.10.617091

**Authors:** Théo Lemaire, Yi Yuan, Celia Gellman, Amy M. LeMessurier, Sarah R. Haiken Dray, Justin P. Little, Robert C. Froemke, Shy Shoham

**Affiliations:** Neuroscience and Tech4Health Institutes and Department of Ophthalmology, New York University Grossman School of Medicine, New York, 10016, USA; Institute of Electrical Engineering, Yanshan University, Qinhuangdao, 066004, China

**Keywords:** ultrasound neuromodulation, two-photon calcium imaging, cortical circuits

## Abstract

Transcranial Ultrasound Stimulation (TUS) can noninvasively and reversibly perturb neuronal activity, but the mechanisms by which ultrasound engages brain circuits to induce functional effects remain unclear. To elucidate these interactions, we applied TUS to the cortex of awake mice and concurrently monitored local neural activity at the acoustic focus with two-photon calcium imaging. We show that TUS evokes highly focal responses in three canonical neuronal populations, with cell-type-specific dose dependencies. Through independent parametric variations, we demonstrate that evoked responses collectively scale with the time-average intensity of the stimulus. Finally, using computational unmixing we propose a physiologically realistic cortical circuit model that predicts TUS-evoked responses as a result of both direct effects and local network interactions. Our results provide a first direct evidence of TUS’s focal effects on cortical activity and shed light on the complex circuit mechanisms underlying these effects, paving the way for TUS’s deployment in clinical settings.

## INTRODUCTION

The brain orchestrates neural activity across a distributed network of specialized neuronal sub-circuits. Exploring the processing of information within and across these circuits necessitates precise tools for causally perturbing neural activity with high spatiotemporal resolution. However, current neuro-manipulation tools are either highly invasive or restricted to superficial brain areas with poor spatial precision^1^.

TUS has recently emerged as a powerful tool for overcoming these limitations. Unlike electromagnetic fields, ultrasonic energy can be focused through the skull into virtually any brain location with millimeter-scale precision^2^. There, it exerts focal mechanical effects on neural tissue that, once integrated over tens to hundreds of milliseconds, can modulate neural activity^3^. At the circuit level, this neuromodulation translates into online functional responses^4,5^ and behavioral effects^6^, but can also induce profound neuroplastic effects that significantly outlast the stimulus^7,8^. Perhaps most interestingly, distinct sonication protocols have been shown to steer neural activity towards either enhanced or suppressed regimes^9,10^. Effectively harnessed, TUS could therefore lead to advanced strategies for the *noninvasive, focal*, and *bidirectional* (i.e., excitatory or inhibitory) control of neural activity, with wide ranging research and translational potential. But which exact stimulation parameters should be used to bias circuits towards a specific activity regime?

Empirical evidence depicts a broad landscape of TUS-evoked effects, with diverse and sometimes conflicting results reported across brain regions, experimental settings, and species^10–12^. However, these effects were mostly examined through the prism of non-specific, aggregated neural readouts^5,7,13^, which generally limits the capacity to dissect the complex interaction between ultrasound and neural circuits. In particular, differences in intrinsic cellular properties (e.g., morphology and ion channel composition)^14^ across neuronal subpopulations are likely to mediate cell-type-specific responses to TUS^15^, which could potentially underlie parameter-dependent transitions between net excitation and suppression of neural activity. Therefore, to better comprehend and predict how ultrasound can manipulate brain activity, we need to evaluate its impact *beyond* the mesoscopic scale, at the level of neuronal populations. Indeed, recent studies have started to address this challenge. In the rodent cortex, electrical recordings have shown that TUS evoked distinct effects in regular and fast spiking units^16^, although this classification does not directly map to cell type and neural function. In subcortical areas, optical recordings have revealed distinct responses in glutamatergic, GABAergic, and dopaminergic neurons^17,18^, albeit only at the level of large aggregated populations through fiber photometry. In addition, these studies used weakly focused ultrasonic beams spanning several millimeters across, thereby inevitably engaging with neighboring brain areas. This lack of selectivity in stimulation, recording, and/or classification poses a significant challenge for the dissection of focal interactions between ultrasound and neural circuits.

To overcome these limitations, we developed a high-resolution bidirectional neural interface combining a two-photon (2P) microscope with a tightly focused ultrasound neuromodulation device. These two systems were integrated in a coaxial arrangement using an annular transducer, allowing us to image through the transducer aperture^19^ while minimizing issues with focal alignment^20,21^, shear waves^22^, reflections and other experimental issues related to ultrasound delivered at an angle or in an unfocused device^12,23,24^. Using this platform, we delivered precise acoustic perturbations to the primary visual cortex (V1) of awake (head-fixed) mice, whilst simultaneously imaging the functional fluorescent activity of specific cortical sub-populations. Specifically, we used different transgenic mouse lines each expressing the calcium sensor GCaMP6s ^25^ in 3 genetically distinct cell types: Thy1-positive pyramidal excitatory neurons, as well as somatostatin-positive (SST) and parvalbumin-positive (PV) inhibitory interneurons (INs). We found that TUS robustly engages cortical neurons to induce functional responses, which are restricted to the acoustic focal area and scale with the time-average intensity of the stimulus. We show that these dose-dependent responses are stereotypical within subpopulations and vary substantially across cell types. Finally, we used a computational model of the V1 microcircuitry to disentangle direct and network-mediated effects of ultrasound on evoked responses.

## RESULTS

### A platform for imaging focal ultrasound neuromodulatory effects at cellular resolution

We developed a combined 2P imaging and ultrasound stimulation platform for use in awake *in vivo* experiments, in order to examine the impact of TUS on local cortical circuits (Figure 1a). To image neural activity at cellular resolution, we coupled our microscope to a high numerical aperture objective focused on the primary visual cortex of awake, head-fixed mice outfitted with a cranial window. This imaging setup produced a 500-by-500 μm imaging plane comprising hundreds of identifiable neurons, allowing for a dense inspection of cortical activity (Figure 1b). To selectively perturb local neural circuits within the optical imaging plane, we used an annular piezoelectric transducer held at a fixed distance from the microscope objective by a 3D-printed holder, allowing for imaging of the murine cortex simultaneously with TUS.

**Figure 1.**
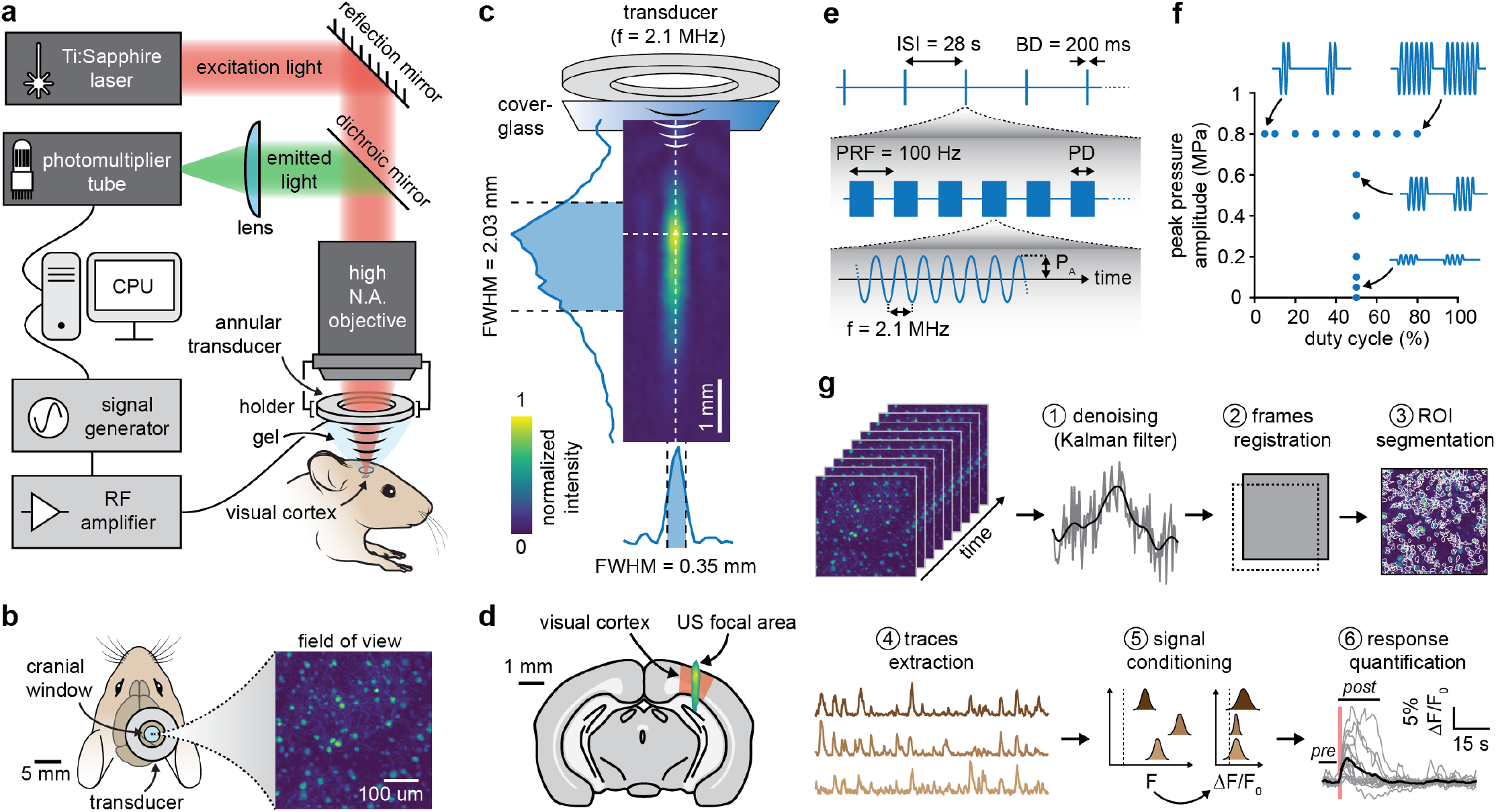
An experimental platform to inspect focal ultrasound neuromodulatory effects in neural circuits. **(a)** Illustration of the experimental setup combining 2P microscopy and ultrasound stimulation. **(b)** Top view of the experimental configuration, depicting the annular transducer, cranial window, and a blow-up of the field of view imaged from the mouse primary visual cortex, comprising hundreds of identifiable neurons. **(c)** 2-dimensional map of the acoustic intensity field generated by the transducer through a cover glass, along with lateral and axial profiles through the acoustic focus. **(d)** Focal transducer intensity field overlaid on an outline of the mouse brain in the coronal plane, highlighting the location of the visual cortex. **(e)** Schematic diagram of the ultrasound stimulation protocol showing the carrier waveform, pulsing parameters, and inter-sonication internal. **(f)** Illustration of the explored values of peak pressure amplitude and stimulus duty cycle during independent experimental trials. **(g)** Schematics of the computational pipeline used to process 2P data and analyze effects of TUS on neural activity including movie denoising and registration, segmentation of cells within the field of view and extraction of their fluorescence traces, conditioning of the traces, aggregation across trials, and response quantification.

To determine which transducer axial position would ensure a robust alignment between the imaging plane and the acoustic focus, we mapped the acoustic intensity field through a glass cranial window, using a scanning hydrophone. Driven at its resonant frequency of 2.1 MHz (Extended Data Figure 1a), the transducer produced an ellipsoidal full width half-maximum (FWHM) focus spanning 0.35 mm laterally (i.e., within the imaging plane dimensions) and 2.03 mm axially, with a peak pressure point located ∼2.1 mm from the radiating surface (Figure 1c). These highly focal properties enable the precise targeting of specific areas of the murine neocortex down to single cortical columns^26^, and are therefore well suited for the causal manipulation of local neural circuits (Figure 1d).

Using this confocal arrangement, we targeted ultrasound perturbations to the primary visual cortex of mice while imaging fluorescence activity in the supra-granular layers (layers 2/3) and within the acoustic focus. TUS stimuli were delivered as short (200 ms long) ultrasound pulse trains with an internal pulse repetition frequency (PRF) of 100 Hz (Figure 1e). Within this conserved pulsing motif, we applied varying levels of peak acoustic pressure amplitude (P_A_, from 0 to 0.8 MPa, see Extended Data Figure 1b) and duty cycle (DC, from 5 to 80%) in two independent explorations (Figure 1f). Finally, we developed an elaborate analysis pipeline to analyze the effects of these perturbations on imaged populations (Figure 1g).

### Ultrasound induces focal responses in cortical neurons

We first assessed whether our platform could reliably capture the response of specific neuronal sub-populations to ultrasound. To this end, we used transgenic mice selectively expressing GCaMP6s in SST-positive cells (Figure 2a). This class of neocortical inhibitory interneurons is found throughout all layers of the neocortex, only projects within the local network, and exhibits a low spontaneous firing activity^27^. As such, it represents a prime target for the observation of direct TUS-evoked cellular responses. Within this population, TUS elicited robust functional responses characterized by a transient increase in fluorescence peaking approximately 1-1.5s after the stimulus onset (Figure 2b,c). Moreover, both the proportion of responding cells and the amplitude of individual cellular responses increased with the amplitude of the acoustic pressure, as reflected in the population average trace. Under sufficiently strong exposure conditions (0.8 MPa, 50 % duty cycle, I_SPTA_ = 10 W/cm^2^), most identified SST cells showed highly robust TUS-evoked responses.

**Figure 2.**
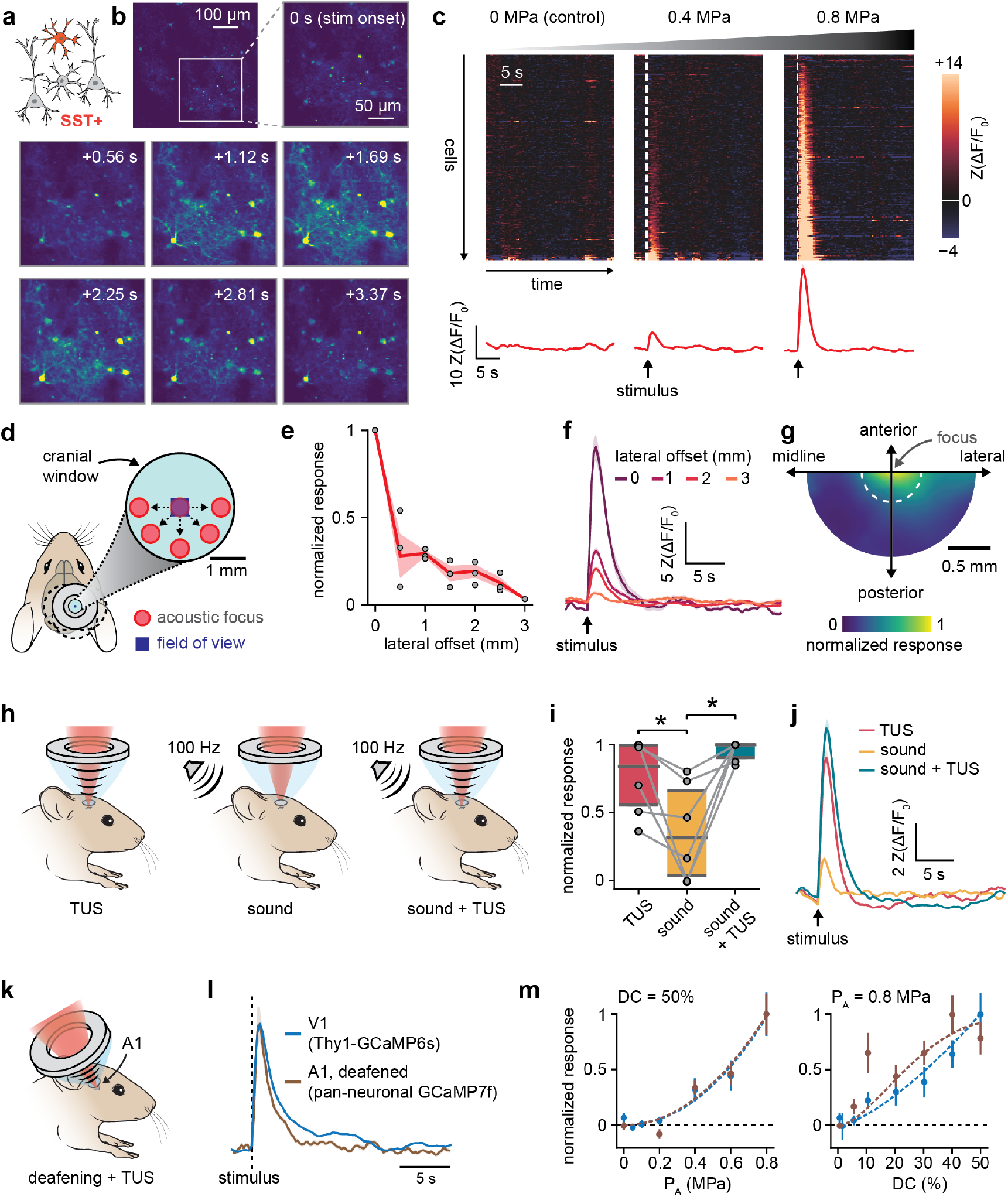
TUS induces focal responses in cortical neurons. **(a)** Illustration of the cell-type-specific imaging of SST-positive cortical cells. **(b)** Representative time lapse images of GCaMP6s fluorescence in a 250-by-250 μm inset of the imaged region before, during and after a TUS stimulus (P_A_ = 0.8 MPa, 50% DC). **(c)** Top: z-scored, and response-magnitude sorted ΔF/F raster plots showing the fluorescence activity evoked by TUS stimuli (200 ms, 50% duty cycle) of varying peak pressure amplitudes in an imaged population of SST INs (n = 234 cells, median of 15 trials). Bottom: z-scored ΔF/F trace (mean +/-SEM) showing the population average response to each pressure level. **(d)** Illustration of the spatial offset control experiment, carried out under strong exposure conditions (0.8 MPa, 50 % duty cycle). **(e)** Normalized amplitudes of TUS-evoked responses in SST INs as a function of the applied lateral offset (n = 3 animals). **(f)** z-scored ΔF/F traces of TUS-evoked responses in SST INs at characteristic offset distances. **(g)** Interpolated heatmap of normalized TUS-evoked response amplitudes the explored 2-dimensional space of lateral offsets (n = 3 animals). The dotted white line indicates the dimensions of the acoustic focus **(h)** Schematics of the buzzing control experiment illustrating the 3 tested conditions; TUS was delivered at 0.8 MPa, 50 % duty cycle. **(i)** Normalized responses amplitudes evoked by each condition across imaged regions. Statistical comparisons between the 3 conditions were performed with a paired t-test, and star symbols indicate significant differences. **(j)** Mean z-scored ΔF/F traces of SST responses evoked in each experimental condition (n = 4 animals). **(k)** Illustration of combined TUS-imaging experiments targeting the primary auditory cortex. **(l)** Normalized mean z-scored ΔF/F traces of TUS-evoked responses in the visual cortex (n = 10 animals, Thy1-specific GCaMP6s reporter, in blue), and in the auditory cortex of a deafened animal (n = 1 animal, pan-neuronal GCaMP7f reporter for identified TUS-responding neurons, in brown); solid lines and shaded areas represent the mean +/-SEM of responses across all selected neurons and for all stimulations with an ISPTA ≥ 2 W/cm^2^). **(m)** TUS-evoked response magnitude in V1 (Thy1-GCaMP6s, in blue) and A1 (pan-neuronal GCaMP7f, for identified TUS-responding neurons, in brown) as a function of peak acoustic pressure (left, DC = 50%) and duty cycle (right, P_A_ = 0.8 MPa); data points depict the mean across neurons, and shaded areas depicted either the weighted SEM across animals (for V1 data, see Figure 3c,e) or the SEM across neurons (for A1 data); dashed lines show polynomial fits to each parameter-dependent response profile.

To examine the focality of these effects, we introduced a lateral offset between the transducer and the objective and measured associated changes in TUS-evoked activity (Figure 2d). We found that response amplitudes decreased monotonically with increasing offset distances, reaching ∼30% of the on-target value at 0.5 mm, and <10% at 3 mm (Figure 2e). This sharp reduction in evoked functional effects closely followed the spatial footprint of the incident pressure field (Figure 1c,d), and was not associated with visible changes in the response temporal dynamics (Figure 2f), nor did it show any visible degree of directionality (Figure 2g). To further verify that observed responses were not confounded by the engagement of peripheral auditory circuits, we performed two control experiments. First, we replaced or complemented the ultrasound stimulus with a buzzing sound at the stimulus PRF (Figure 2h). We found that while the sound stimulus evoked responses in SST INs, their amplitude was significantly lower than that evoked by TUS (p = 0.039, paired t-test) or by a combination of TUS and sound (p = 0.003, paired t-test) (Figure 2i,j). Second, we targeted our platform to the mouse primary auditory cortex (A1) and examined TUS’s effects on local neurons using a pan-neuronal fluorescent reporter (Figure 2k and Extended Data Figure 2a). We also observed robust responses, albeit with a longer duration and slower dynamic than those observed in V1, possibly indicating the involvement of both direct and indirect effects (Extended Data Figure 2b). To disentangle these effects, we performed a bilateral surgical deafening procedure that effectively abolished auditory brainstem responses (Extended Data Figure 2c). Following this intervention, we observed strong, unequivocally direct TUS-evoked responses in A1, with a temporal dynamic and parameter dependency closely resembling those seen in V1 (Figure 2l,m and Extended Data Figure 2b,d). Collectively, these results provide strong evidence that TUS causally and directly excites cortical neurons through the focal action of the acoustic energy.

### Ultrasound-evoked responses are cell-type-specific, stereotyped, and dose-dependent

To carry out efficient computations, the brain relies on a diverse set of neuronal populations with distinct functional specializations. In the primary visual cortex, the majority of neurons (>90%) can be classified into one of 3 non-overlapping cell types: Thy1 excitatory neurons, SST INs, and PV INs^28^. We compared the responses evoked by various TUS stimuli in these three populations (Figure 3a), to examine their hypothesized cell-type specificity. To narrow down our parametric exploration, we specifically examined the impact of two critical parameters – the peak pressure amplitude and the stimulus duty cycle – because of the abundant evidence supporting their crucial role for ultrasonic neuromodulation^29^.

**Figure 3.**
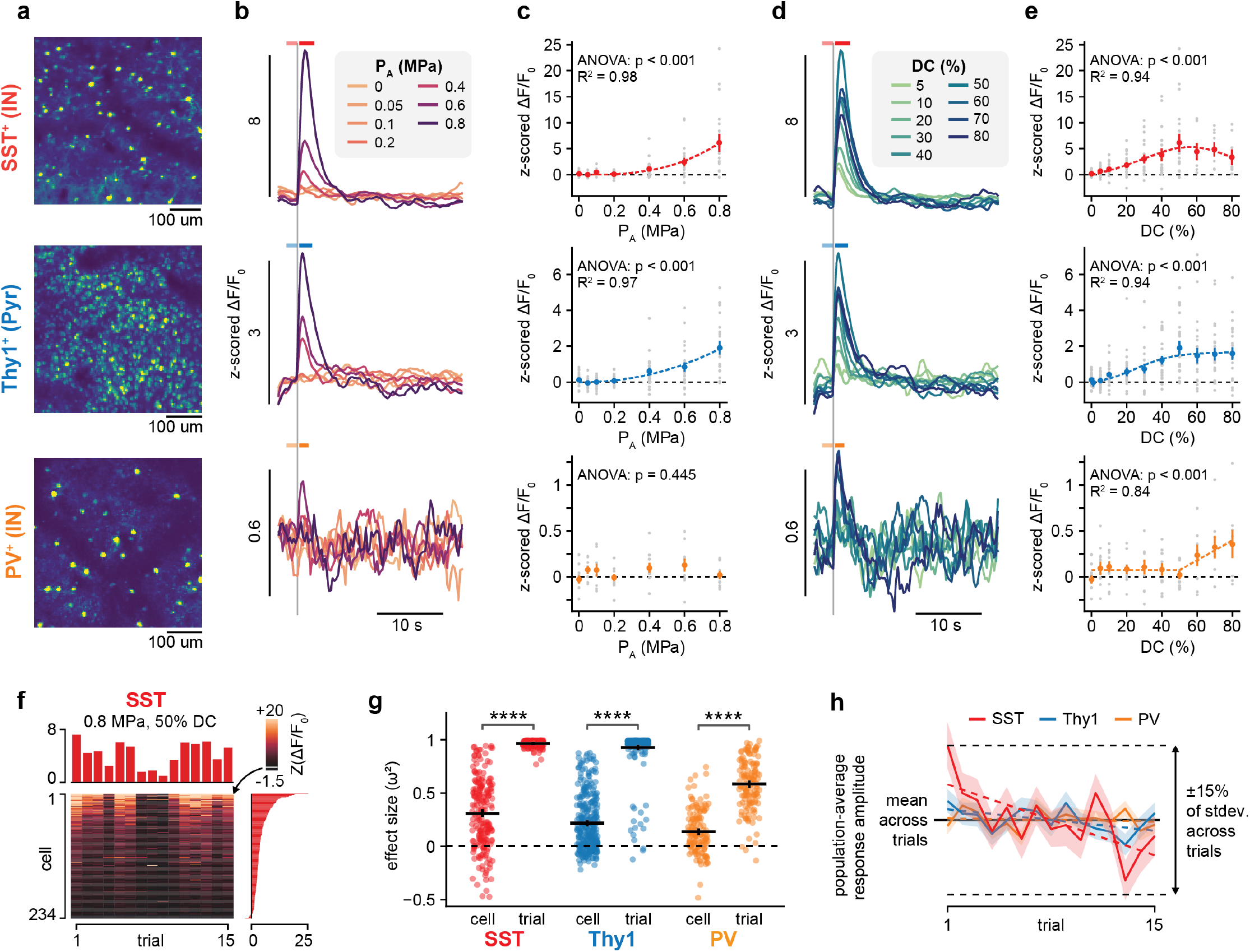
TUS induces cell-type-specific and dose-dependent responses in cortical neurons. **(a)** Maximal projection images (saturated at the 99^th^ percentile) of representative fields of view illustrating the distribution of the three imaged neuronal populations. **(b)** Mean z-scored ΔF/F traces of TUS-evoked responses in SST+ (top, n=6 animals), Thy1+ (middle, n=10) and PV+ (bottom, n=7) neurons for distinct peak acoustic pressures at constant duty cycle (DC = 50%). Light and dark colored horizontal bars respectively indicate the pre- and post-stimulus windows used for the response quantification shown in (c). **(c)** TUS-evoked response magnitude as a function of peak acoustic pressure in the same 3 populations; grey data points denote average response magnitudes for each imaged region; colored data points and error bars depict the mean +/-SEM across animals, weighted by the number of imaged neurons in each animal; R^2^ values indicate the accuracy of best fits (dashed lines) for populations in which parameter dependency was confirmed by 1-way ANOVA. **(d, e)** Mean z-scored ΔF/F traces (d) and TUS-evoked response magnitude (e) as a function of stimulus duty cycle at constant pressure amplitude (P = 0.8 MPa) in the same 3 populations. **(f)** Exemplar heatmap of evoked response magnitudes from a population of SST INs for 15 consecutive stimulus presentations at 0.8 MPa, 50% DC. Marginal plots show the population average response to each trial (top) and the trial average response of each ROI (right). **(g)** Distribution of effect sizes (ω^2^) computed from the results of 2-way (cell, trial) ANOVAs performed independently across animals and stimulation conditions. Individual points denote effect size for one animal and condition; horizontal and vertical black bars denote the mean +/-SEM of each distribution. Stars indicate statistically significant differences between the “trial” and “cell” effect size distributions, for each neuron type (Mann-Whitney test, ****: p < 10^−4^). **(h)** Z-scored vectors of population-averaged response magnitudes across trial sequences. Solid colored lines and shaded areas indicate the mean +/-SEM of z-scored response vectors, while dashed colored lines indicate fitted linear regressors for each neuron type. The solid black line represents the mean response across trials (at which all trial vectors were aligned), while dashed horizontal black lines indicate +/-15% of the standard deviation across the trial sequence (with which all vectors were normalized).

We first probed the effects of pressure amplitude variations at a fixed duty cycle (DC = 50%). Both Thy1 and SST populations showed clear pressure dependencies, with a response threshold around 0.2 MPa followed by a monotonic increase in response magnitude with increasing pressure levels (Figure 3b,c). These dependencies were accurately captured by quadratic fits (R^2^ = 0.98 and 0.97 for SST and Thy1 neurons respectively). In contrast, the PV population did not exhibit a significant dependency on pressure (p = 0.445, 1-way ANOVA).

We then probed the effects of the stimulus duty cycle at a fixed peak pressure amplitude (P_A_ = 0.8 MPa) and observed more complex dependencies across neural sub-populations (Figure 3d,e). Both SST and Thy1 neurons responded at very low DC and their response magnitudes gradually increased with larger DCs, reaching a maximum for DC = 50%; at higher DCs these increases saturated, and the SST responses even weakly decreased. In contrast, PV neurons did not respond at low duty cycles, and showed a gradual increase in response magnitudes at higher doses (DC > 50%). These complex dependencies were best captured by distinct analytical functions: a sigmoidal fit for Thy1 neurons (R^2^ = 0.94), a combination of sigmoidal increase and exponential decay for SST neurons (R^2^ = 0.94), and a threshold-linear function for PV neurons (R^2^ = 0.84). The cell-type-specific nature of these responses and their parameter dependencies – particularly the pronounced differences in activation thresholds across cell types – strongly suggests that different cortical sub-populations possess distinct intrinsic sensitivities to ultrasound. Moreover, the existence of intricate, non-monotonic dependencies supports the hypothesis that TUS-evoked responses are actively modulated by local network projections.

Beyond their cell-type-specific parameter dependencies, we questioned whether – and to which extent – TUS-evoked responses would vary across cells inside a population. For all 3 neuron types, the relative response strengths of individual cells were strongly conserved across trials, despite substantial variations in the population’s average response (Figure 3f). Indeed, a 2-way analysis of variance (ANOVA) revealed much larger effect sizes across trials than across constituent cells, irrespectively of cell type and stimulation parameters (Figure 3g). Moreover, distributions of trial-to-trial response amplitudes between co-registered cells were significantly positively correlated, and a substantial portion of the response variance across trials could be projected along a single vector (Extended Data Figure 3a,b). Given the high population stereotypy of responses, we further examined response variation across trials on population averages. After z-scoring each response vector to a normalized space, we observed that most responses followed a unimodal normal distribution (Extended Data Figure 4a-c). Stationarity analysis revealed a slight decrease in response magnitude throughout trial sequences (Figure 3h), but this decrease was only significant in SST cells, where the average dip per trial amounted to less than 1% of the response standard deviation along the trial sequence (Extended Data Figure 4d). Collectively, these results indicate that TUS evokes highly stereotyped population responses with no evidence of fading or cumulative effects, irrespectively of cell type and stimulation parameters.

### Ultrasound-evoked responses scale with the time-average acoustic intensity

Beyond the pronounced differences observed across sub-populations, our parameter exploration revealed that cortical neurons show distinct dependencies on the stimulus pressure amplitude and duty cycle. We therefore wondered whether we could identify a unifying scaling law that would capture the neuromodulatory effect of these two independent parameters. We hypothesized that if TUS-evoked responses scale with an acoustic dose function *γ* = (*P*_*A*_, *DC*), response curves obtained from our pressure and duty cycle sweeps would naturally align when projected onto that dose space (Figure 4a). Because pressure amplitude and duty cycle are intrinsically orthogonal parameters (P_A_ scales the stimulus “vertically” while DC acts on its temporal dimension), we conjectured that their compound effect would be multiplicative, rather than additive. Therefore, we considered 2 customary measures of acoustic dose that each multiply pressure and duty cycle with distinct scaling effects (Figure 4b and supplementary information): the spatial-peak, time-average peak pressure amplitude (*P*_*SPTA*_ = *P* · *DC*), and the spatial-peak, time-average acoustic intensity 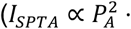.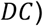. To complement our analysis, we considered a third measure with a nonlinear dependency on duty cycle: the spatial-peak, root-mean-square acoustic intensity 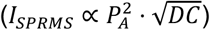.

**Figure 4.**
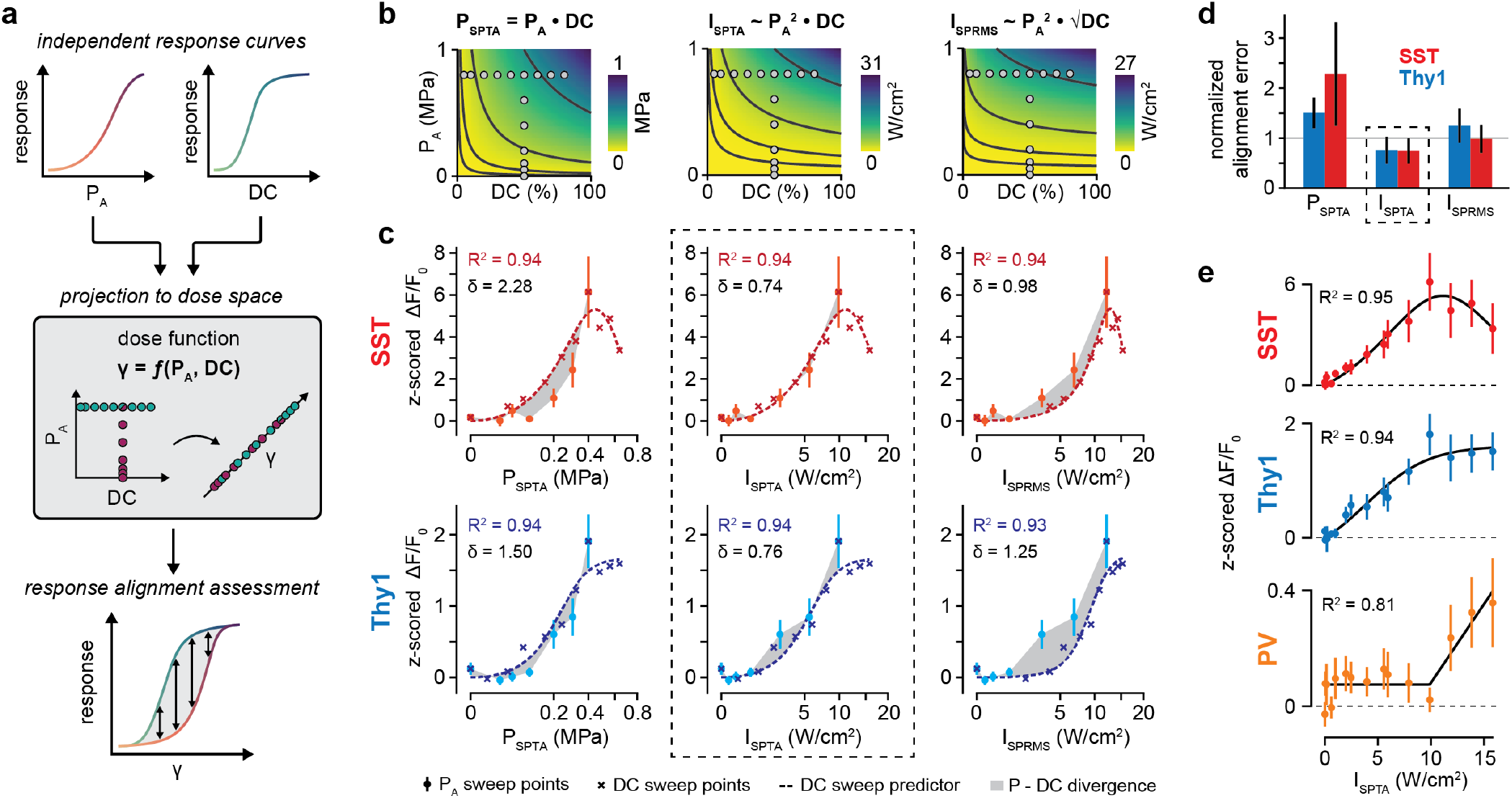
TUS-evoked neural responses scale with the time-average acoustic intensity. **(a)** Schematic diagram illustrating the concept of dose projection to evaluate compound neuromodulatory effects. **(b)** Heatmaps of examined acoustic dose functions across the 2-dimensional P_A_-DC space (left: P_SPTA_, middle: I_SPTA_, right: I_SPRMS_). Grey points represent (P_A_, DC) combinations explored in parametric sweeps; black lines represent characteristic dose levels logarithmically distributed at 0.5, 2.3, 10.7 and 50% of the maximal dose value. **(c)** Projected response curves of the pressure amplitude and duty cycle sweeps as a function of the 3 examined acoustic dose functions in SST+ neurons (top, in red) and Thy1+ neurons (bottom, in blue). Horizontal axes are presented on a square root scale to facilitate the visualization of the response regimes. Light-colored points and error bars depict the weighted mean +/-SEM of data points in the pressure amplitude sweep. Dark colored crosses indicate the weighted mean of data points from the duty cycle sweep, and dashed lines represent the fitted predictor function to the projected DC response curves. R-squared values indicate the goodness-of-fit of DC predictor functions for each dose projection. Grayed areas indicate the deviation between data points from the pressure amplitude sweep and interpolated points from the DC fitted predictor, for which mean SEM-normalized alignment errors are also indicated. The dashed rectangle highlights projections with the best alignment. **(d)** SEM-normalized alignment error between data points from the pressure amplitude sweep and interpolated points from the DC fitted predictor at the corresponding dose levels. Bars and error bars indicate the mean +/-SEM of normalized errors for each condition. The gray line materializes the SEM reference level. The dashed rectangle highlights projections with the lowest alignment error. **(e)** Normalized response curves projected in the I_SPTA_ space for all 3 sub-populations. R-squared values indicate the goodness-of-fit of global predictors (black lines).

For each candidate, we inspected the alignment of projected response curves in SST and Thy1 neurons, as these populations were found to be sensitive to both input parameters. Under the P_SPTA_ projection, response magnitudes extracted from the pressure amplitude sweep were lower than those predicted from the DC sweep, for both populations (Figure 4c). In contrast, under the I_SPRMS_ projection, response strengths from the pressure amplitude sweep consistently exceeded those predicted from the DC sweep. In between these two “extreme” scenarios, the I_SPTA_ projection produced a robust alignment between our two parametric sweeps, consistently across the dose range. To confirm these qualitative trends, we computed for each candidate projection the scatter-normalized absolute error (δ, see Methods) between (1) projected data points from the pressure amplitude sweep and (2) a fitted predictor of the projected DC response curve interpolated at the corresponding dose levels. Expectedly, the I_SPTA_ projection consistently yielded the lowest alignment error across both Thy1 and SST populations, with deviations significantly lower than the intrinsic response SEM (Figure 4d). This result was not influenced by the quality of DC response predictors, as DC fitting performance was highly accurate across all 3 projections (Figure 4c).

Re-fitting our predictor functions on aggregated data in the I_SPTA_ space, we obtained remarkably accurate analytical predictions of dose-dependent response scaling in all populations, including PV neurons (Figure 4e). In this one-dimensional dose space, a clear phenomenological bifurcation emerges. Low intensities (I_SPTA_ ≤ 10 W/cm^2^) induce gradually increasing responses in both SST and Thy1 neurons while PV neurons stay predominantly quiet. In contrast, high intensities (I_SPTA_ > 10 W/cm^2^) lead to a sharp increase in PV responses accompanied by a decrease in SST responses and a saturation of Thy1 responses.

Beyond their robust dose scaling, we wondered if TUS-evoked responses could also be modulated by the ongoing neural activity at the time of stimulation, as is the case with electrical stimulation^30^. Binning population-average traces by their pre-stimulus activity levels revealed a clear anti-correlation between pre-stimulus activity and post-stimulus activity change, for all 3 cell types (Extended Data Figure 5a,b). In particular, the decrease in post-stimulus activity seen for elevated pre-stimulus levels suggests a potential TUS inhibitory effect when circuits are strongly active. However, we found similar anti-correlation levels on surrogate datasets lacking a stimulus-evoked component (Extended Data Figure 5c,d), suggesting that observed dependencies might simply arise from the autocorrelative nature of our data. Therefore, to assess the “true” dependency of TUS-evoked responses on ongoing neural activity, we subtracted “expected” post-stimulus trends from our signals using autoregressive (AR) models. These AR-corrected responses showed much weaker anti-correlations with pre-stimulus activity, and did not undergo any stimulus-induced decrease in activity even when pre-stimulus activity was high (Extended Data Figure 5e,f). These results indicate that, rather than having an activity-dependent bidirectional (i.e., excitatory or inhibitory) effect on neural activity, TUS is simply less effective at exciting circuits that are already strongly engaged.

### A self-sufficient and physiologically realistic mechanism for TUS-evoked cortical circuit responses

To process sensory inputs, the primary visual cortex relies on a well-established microcircuit architecture^28,31^. In this circuit, pyramidal neurons form excitatory projections onto both SST and PV interneurons, which in turn provide feedback inhibition to the distal dendrites and peri-somatic regions of pyramidal neurons, respectively. This intricate functional network regulates neural activity at speeds approaching the millisecond timescale and is therefore likely to influence the response of neurons to an incident TUS perturbation. As such, it also confounds the analysis of direct TUS-neuron interactions and their potential cell-type-specificity. For example, the decrease in the responses of SST INs at high intensities could be an intrinsic property of TUS-SST interactions, or (more likely) result from their synaptic inhibition by PV cells.

Therefore, to infer the relative contributions of direct and network-mediated effects in observed TUS-evoked responses, we developed a computational model of the neocortical microcircuitry incorporating our three interconnected populations (Figure 5a). This model describes the time-varying activity of each neural population as a function of their current activity state and the transformation of their physiological inputs (i.e., pre-synaptic and stimulus-mediated) by an activation function (see Methods). To mimic the excitability and response properties of cortical neurons, we modeled these activation functions as rectified linear units with specific threshold (*θ*) and gain (*g*) parameters^32^. To tailor the model to our experimental context, we used publicly available physiological datasets from the Allen Brain Atlas (see Methods) to derive a reference set of gain parameters and coupling weights describing the specific cellular properties of excitatory, SST, and PV neurons and their interconnectivity in the V1 micro-circuitry (Figure 5b,c). To model TUS’s impact on the system, we added a time-varying input representing the incident ultrasound stimulus, converted to a physiological current via multiplication by a cell-type-specific sensitivity term (λ). We used a multi-dimensional genetic algorithm optimization to assess whether a specific set of model parameters (i.e., stimulus sensitivities and network architecture) could reproduce the empirically observed response profiles, while minimizing deviations from the reference coupling weights (see Methods and Extended Data Figure 6). Across multiple separate runs with random initializations, the optimizer converged towards a highly consistent set of model parameters (Figure 5d,e) that accurately reproduced our empirical response profiles across all 3 populations (Figure 5f). Notably, this optimization required only marginal adjustments to the model’s original connectivity matrix from the Allen Brain Atlas *Synaptic Physiology dataset* (average absolute relative deviation in coupling weight of 4.6 ± 2.5%, range 0-18%), implying that physiological network architectures can explain our data. Complementarily, the optimizer revealed substantial (and statistically significant) differences in intrinsic ultrasound sensitivity across our populations (Figure 5e): excitatory neurons emerged as the most sensitive, followed closely by PV neurons (*λ*_*PV*_ ≈ 0.87 *λ*_*E*_), while SST neurons exhibited a much lower relative sensitivity (*λ*_*SST*_ ≈ 0.15 *λ*_*E*_). Notably, when constrained by imposing a uniform stimulus sensitivity, the optimizer converged towards a nonrealistic network architecture and failed to replicate fundamental features of our empirical response profiles yielding a 5-fold increase in prediction errors (Extended Data Figure 7a-d), implying that differential sensitivity is crucial to the accuracy of our predictions.

**Figure 5.**
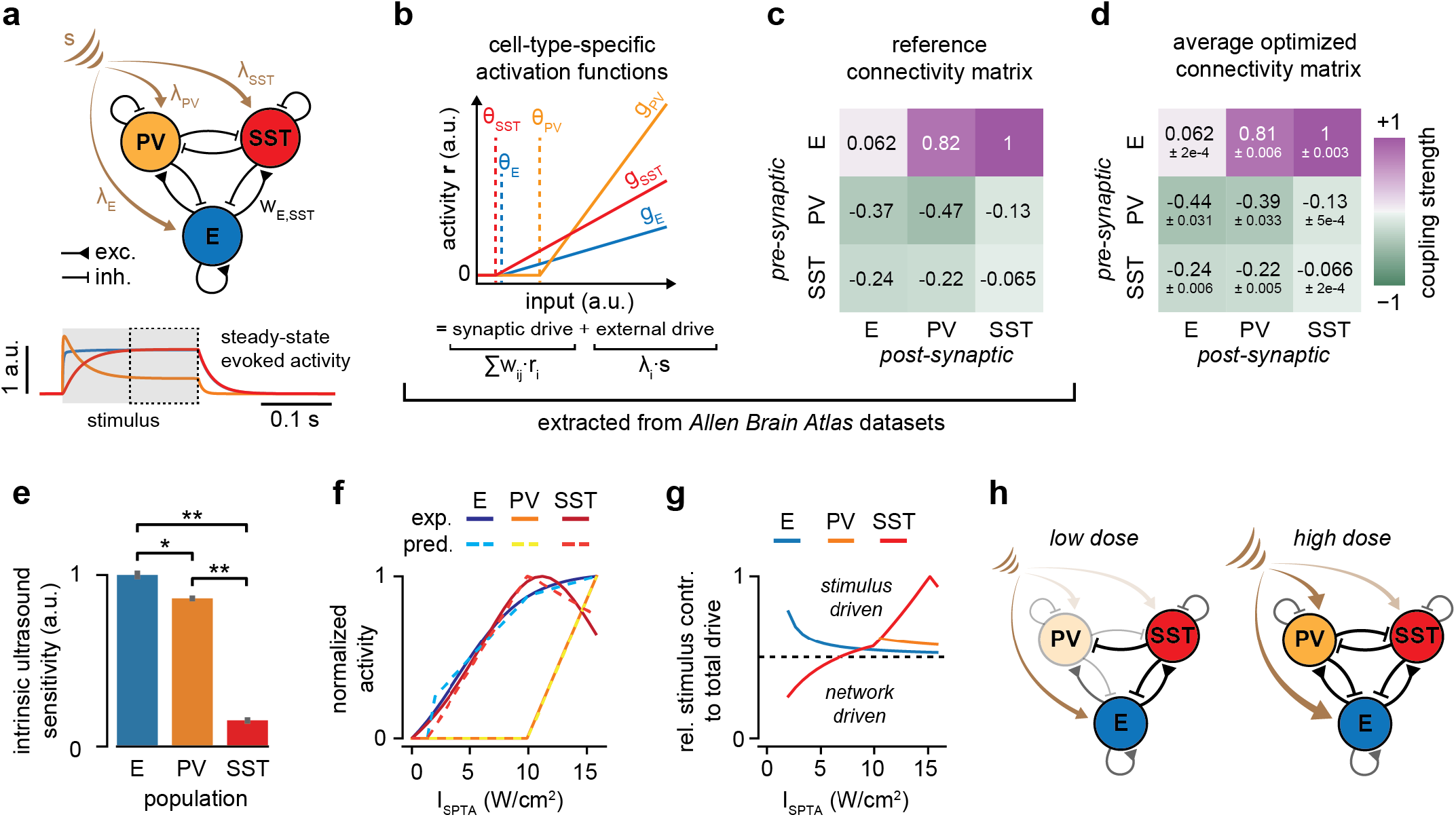
A cortical microcircuit model predicts that TUS-evoked responses require cell-type-specific ultrasound sensitivities. **(a)** Top: schematic diagram of the V1 microcircuit model supplemented with TUS-dependent inputs (brown arrows). Bottom: exemplar model simulation output predicting time-varying, TUS-evoked activity profiles in excitatory, PV and SST neurons. The shaded area denotes the stimulus interval; the dashed area indicates the steady-states extraction window. **(b)** Cell-type-specific activation functions derived from the *Cell Feature* dataset of the Allen Brain Atlas. Dashed vertical lines represent cell-type-specific activation thresholds in the model input space. **(c)** Reference connectivity matrix of the mouse V1 micro-circuitry extracted from the *Synaptic Physiology* dataset of the Allen Brain Atlas. Excitatory (inhibitory) connections are depicted in purple (green), with a color saturation proportional to the coupling weight magnitude. **(d)** Optimal connectivity matrix obtained across 5 model optimization runs. Mean +/-std values are indicated for each coupling weight. **(e)** Distribution of optimal cell-type-specific stimulus sensitivities obtained across 5 model optimization runs. Bars and error bars depict the mean +/-SEM of sensitivities normalized to the unit interval. Stars denote statistically significant differences between distributions (Mann-Whitney test, *: p < 0.05, **: p < 0.01). **(f)** Comparison of cell-type-specific and I_SPTA_-dependent response profiles predicted by the average optimized model against their empirical counterparts. Solid lines denote the empirically fitted predictors, while dashed lines denote the model predictions. Profiles were normalized to the unit interval. **(g)** Proportion of the total cellular input provided by the stimulus in evoked responses across the I_SPTA_ range for each population. The dashed horizontal represents the 50% mark where both network and stimulus contribute equally to cellular inputs. **(h)** Conceptual representation of the predicted circuit mechanisms underlying TUS-evoked responses at both low (left) and high (right) ultrasonic doses.

Beyond replicating our experimental results, this computational deconstruction effectively unpacks TUS-evoked circuit responses into separate, quantifiable contributions from the stimulus and from network interactions (Figure 5g). This deconstruction delineates two clear “circuit regimes” (Figure 5h) that mirror the dose-dependent phenomenological bifurcation seen in our empirical data (Extended Data Figure 8a,b): at low doses, TUS directly activates excitatory neurons; their activity preferentially recruits SST neurons, which in turn regulate overall excitation and keep PV neurons silent. At higher doses, the increased drive from both stimulus and pyramidal activity recruits PV neurons, which in turn start inhibiting SST neurons and therefore gradually take over as the main circuit regulator.

## DISCUSSION

Transcranial Ultrasound Stimulation emerges as a promising noninvasive clinical solution to treat a variety of neurological conditions including essential tremor, epilepsy, Parkinson’s disease, depression, drug abuse disorder, and schizophrenia^33,34^. Yet, TUS is still fundamentally hindered by a lack of understanding of its underlying mechanistic principles. To address this gap we conducted here an in-depth characterization of TUS’s neuromodulatory effects in the mammalian neocortex. We show that ultrasound exerts a focal action on neural tissue which induces responses in both excitatory and inhibitory cortical neurons, at exposure levels that have been safely used in non-human primates^5,6^ and human subjects^33^. These responses show clear dependencies on two independent parameters (pressure amplitude and duty cycle), which can be unified under a common scaling law based on the time-average acoustic intensity of the stimulus. Using a tailored computational model, we predict that TUS-evoked responses are best explained by cell-type-specific sensitivities to ultrasound, producing intrinsic activity that is actively modulated by local network projections.

Characterizing TUS’s direct neurophysiological effects and their focality has been a longstanding challenge for the field. Electrophysiology and fiber photometry readouts, which have often been paired with TUS, require the problematic implantation of solid elements at/above the region of interest that may vibrate upon sonication, leading to indirect neural activation. Additionally, these methods provide little spatial information about evoked effects. Noninvasive alternatives such as widefield calcium imaging also require careful consideration. In fact, two prominent TUS studies in rodents reported TUS-evoked cortical responses extending far beyond the putative acoustic focus^12,23^. However, both employed sub-MHz carrier frequencies typically used in human studies, producing ultrasonic beams wide enough to irradiate large portions of the rodent brain and reflect off the skull’s base. In addition, by delivering ultrasound obliquely, they favored the generation of shear waves that can propagate across the skull to affect distant areas^35^. Here, we show that delivering tightly focused 2.1 MHz ultrasound perturbations normally to the brain surface elicits highly focal functional responses in a murine brain. Awake preparations seem to be of particular importance in this context, as parallel investigations in anesthetized mice did not reveal TUS-evoked cortical calcium responses at similar dose levels^36^. Laterally, our responses were largely confined to the acoustic focal area (∼0.5 mm radius) with some residual activation spreading up to a few millimeters, likely due to strong lateral coupling of cortical circuits, as seen in focal optogenetic manipulations^37^. Axially, our transducer calibration indicates that the acoustic field likely engaged neural circuits across the entire depth of the murine cortex. Focused transducers or acoustic lenses could help in restricting this axial focus; however, technological advances are needed to integrate such approaches with coaxial high-precision imaging. Alternatively, increasing carrier frequencies could further improve spatial resolution; however, TUS-evoked effects reported in this regime seem primarily inhibitory^20^, suggesting a different – likely thermal – interaction mechanism. Our experimental setup thus represents an attractive trade-off between stimulation focality and clinical relevance.

Aside from interface considerations, the vast array of adjustable TUS parameters implies a critical need to carefully characterize their (co-)influence on TUS’s evoked effects. Here, we showed that independent variations in pressure and duty cycle can be unified under a common scaling law based on time-average intensity. This finding has strong implications for stimulus design, whereby altering duty cycle by a factor *x* is equivalent to altering pressure by a factor 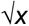.It also hints at an energy transduction mechanism dependent on the integral of the stimulus over time^38^, rather than influenced by discrete ON-OFF transitions. As such, it suggests that excitation is proportional to stimulus duration^3,9^, but contrasts with observed dependences on pulse repetition frequency at constant I_SPTA_ levels^16,39^. This putative discrepancy could indicate limits to the applicability of this scaling law, or alternatively result from methodological differences, e.g., in sonication strategy: delivering ultrasound at an angle^16^ or diffusely through the jaw^21,40^, or the use of anesthesia. Further work is thus needed to clarify possible parametric inter-dependencies and formulate a truly unifying scaling law governing TUS-evoked effects across the entire parameter space. Nonetheless, our I_SPTA_ scaling already enables a comparative re-interpretation of TUS-evoked effects across studies. In particular, the order-of-magnitude difference in the activation thresholds of excitatory and PV inhibitory neurons aligns with fiber photometry observations in the mouse hippocampus^17^, and corroborates electrophysiological results in the mouse somatosensory cortex where low pulsing rates (i.e., low I_SPTA_) evoked activity in regular spiking (i.e., putative excitatory and SST) neurons but not in fast-spiking (i.e., putative PV) neurons^16^. Putatively, this low PV responsiveness arises from their specific electrophysiological properties (a low input resistance leading to a high intrinsic activation threshold, see Figure 5b) rather than a distinctly low ultrasound sensitivity (Figure 5e).

Beyond stimulus design, TUS’s maturation is also hindered by the intrinsic complexity of neural circuits. Most brain areas are comprised of a variety of neuronal populations arranged in intricate functional networks that modulate TUS-evoked responses. Deconstructing which effects are directly induced by ultrasound and how they are integrated into a high-level circuit response would effectively enhance results’ interpretability and their transferability across brain areas. The slow kinetics of our GCaMP6s calcium indicator (> 100 ms) precluded a latency-based deconstruction. Hence, we turned to a computational approach leveraging physiologically realistic cellular and network properties from the Allen Brain Atlas *Synaptic Physiology* dataset^31^, incorporated into a Wilson-Cowan model of excitatory-inhibitory interactions^41^ extended to our 3 populations of interest^42^. Using this microcircuit model, we showed that our empirically observed TUS-evoked responses could naturally arise from the local network when the stimulus is converted into physiological inputs via cell-type-specific ultrasound sensitivities. Future investigations involving the selective inhibition of spiking activity and/or synaptic transmission, as well as faster calcium^43^ or voltage indicators, will be needed to confirm these predictions. Nonetheless, our computational deconstruction delineates a self-sufficient, robust, and physiologically plausible mechanism underlying TUS-evoked responses in cortical circuits.

As it starts to enter clinical settings, TUS critically needs robust mechanistic insights to ensure this technology both safe and effective. Here, we provide strong evidence for TUS’s focal direct effects on cortical activity, and establish a powerful and generalizable framework to understand and predict TUS-evoked effects on the mammalian brain as a function of dose and network composition. This framework could inform the design of effective stimulation protocols as a function of targeted areas and desired functional outcomes. Our findings thus advance TUS’s applicability as a high-precision noninvasive brain perturbation technique for both clinical applications and fundamental neuroscience research.

## MATERIALS AND METHODS

### Mouse lines

Transgenic mice expressing GCaMP6s in pyramidal neurons were obtained from the Thy1.2-GCaMP6s founder line 3, already described in previous work^44^. PV^IRES-Cre 45^ (Stock No. 017320, Jackson Laboratories) and SST^IRES-Cre 46^ (Stock No. 018973, Jackson Laboratories) mice were crossed with Cre-dependent GCaMP6s reporter mice (TIT2L-GC6s-ICL-TPT, Ai163 line) ^47^ to obtain transgenic mice with PV and SST-specific GCaMP6s expression. For auditory cortex experiments, we used SST^IRES-Cre^ mice crossed with Cre-dependent TdTomato reporter mice ^48^ (Ai9, Stock No. 007909, Jackson Laboratories). Male and female mice between 2 and 4 months old were used in all experiments and handled in accordance with institutional guidelines. All procedures were approved under the New York University Langone Health institutional animal care and use committee (IACUC) protocol IA17-01720.

### Surgical preparation

Mice were anesthetized with isoflurane flowing at 1.5 L / min (3.0% during induction, 1.5% during surgery). After securing the head in a stereotaxic frame and applying an ophthalmic ointment, the hair on top of the skull was removed, and the area was sterilized. A longitudinal incision was made to expose the entire dorsal surface of the skull. The periosteum was removed while leaving the bone intact. A circular craniotomy with a diameter of 3 mm was performed over the right hemisphere (centered 2.4 mm lateral to the midline and 2.8 mm posterior from bregma), using an air-driven dental drill (Midwest Tradition, FG 1/8 drill bit), to expose the visual cortex. For auditory cortex experiments, an equivalent craniotomy was performed centered 4 mm lateral to the midline and 3 mm posterior from bregma to expose the auditory cortex, and an adeno-associated viral (AAV) vector encoding the calcium indicator GCaMP7f (pGP-AAV-syn-jGCaMP7f-WPRE, Plasmid #104488, Addgene) was injected using a stereotactic syringe pump (World Precision Instruments Inc.) at a rate of 0.1 μL/min (500 nL total, 300-500 μm deep). A cranial window was then implanted replacing a circular piece of skull by a two-layer glass coverslip (Warner Instruments, USA) that was secured in place using a mix of self-curing resin (Orthojet, Lang Dental, USA) and cyanoacrylate glue (Krazy Glue, USA). Two steel rods were attached to the rostral and caudal extremities of the skull using C&B Metabond dental cement (Parkell, USA). Each animal recovered for at least 10 days prior to experiments.

### Surgical deafening

Mice were anesthetized with isoflurane (see above). A postauricular incision was made and the superficial fascia of the neck was dissected. The sternocleidomastoid muscle and posterior belly of the digastric muscle were retracted from the tympanic bulla to expose the round window, and a bullostomy was performed using a dental drill with a 0.5 mm burr. A sham cochlear implant (CI) array designed for rodents (4-channel, Cochlear) was used for the deafening procedure due to its flexibility that facilitated insertion into the cochlea. The sham CI was inserted into the cochlea to a depth of ∼2.25 mm and left for 10 min to induce full field deafness. The array was removed, and a graft of fascia and muscle was inserted around the round window to prevent leakage of perilymph. The graft was secured with 2-octyl cyanoacrylate (Surgi-Lock 2oc, Meridian Animal Health). The surgical incision was closed with absorbable suture (Ethicon, USA). The procedure was then repeated on the contralateral side.

### Acoustically evoked auditory brain stem responses

Auditory brain stem responses (ABRs) to acoustic tones were assessed before the deafening procedure and 2 weeks post-operatively. Animals were anesthetized with intramuscular ketamine (80 mg/kg) and xylazine (16 mg/kg) and body temperature maintained. Subdermal needle electrodes were placed at the cranial vertex (recording electrode), behind each pinna, and at the base of the spine above the tail (reference and grounds). ABRs were recorded with a preamplifier (DAM50; World Precision Instruments, USA) connected to an amplifier (MultiClamp 700A; Molecular Devices, USA) and digitizer (Digidata 1440A; Molecular Devices, USA). Acoustic tone stimuli (50 ms long, 5 ms rise/fall, 16 kHz carrier frequency, 90 dB sound-pressure level) were presented with a digital signal processor and calibrated speaker (Tucker-Davis Technologies, USA) at a rate of 16.7 Hz for 512 sweeps. Stimulus and ABR waveforms were acquired using Clampex 10.7 (Molecular Devices, USA).

### Transducer assembly and calibration

A micro coaxial cable (9432 WH033, AlphaWire, USA) was stripped at both extremities. Exposed wires on one end were soldered on each side of a lead zirconate titanate (PZT-4) piezoelectric ceramic ring transducer with 1 mm thickness, ID 5mm and OD 10mm (Beijing C.C.W. Ultrasonic Science & Technology, China), giving a transverse resonance mode of 2.1 MHz. On the other end, wires were soldered to a BNC connector. The ring transducer was then glued inside a custom 3D printed holder featuring a dedicated 10 mm diameter circular lip, and a hollow base designed to fit the microscope objective.

Assembled transducers were calibrated inside a custom 3D printed tank filled with degassed deionized water, both in free-field conditions and through a glass window replicating the experimental conditions. The transducer was driven by a waveform generator (DG 1022Z, Rigol Technologies, USA) connected to a 30W, 40 dB voltage amplifier (LZY-22+, Mini-Circuits, USA). Generated acoustic signals were measured by a needle hydrophone (NH-0040, Precision Acoustics, UK) connected to a fast-sampling digital oscilloscope (Model 2555, B&K Precision, USA) via a pre-amplifier, a DC coupler, a 25 dB booster amplifier, and a 50 Ω BNC feed-through terminator. Digitized signals were mean-corrected, band-passed filtered between 1 and 15 MHz to remove electrical noise, and their envelope was extracted by Hilbert transformation to quantify the peak pressure amplitude.

To map the pressure field, we scanned a 3D rectilinear grid spanning 6 mm in the transducer axial dimension and 2 mm across both transverse dimensions with a granularity of 0.1 mm, using an automated micro-manipulation system (MP-285A, Sutter Instruments). Once the location of the focus was established, we measured the peak pressure amplitude for a range of input voltages between 0 and 1.3 Vpp to map the transducer input-output curves.

### Coupled two-photon imaging and ultrasound stimulation

Imaging was performed on an Ultima II multiphoton microscope (Bruker, USA) or a Bergamo II (Thorlabs, USA) system. Two-photon excitation light was emitted at 920 nm using a mode locked, 80 MHz repetition rate, femtosecond-pulsed laser with dispersion compensation (InSightX3, Spectra-Physics, USA, or Chameleon discovery, Coherent, USA). The beam was relayed and magnified by a telescope to the back-aperture of a 10x water immersion objective lens with either 0.6-NA, 8 mm working distance (XLPLN10XSVMP, Olympus, USA) or / 0.5-NA, 7.77 mm working distance (TL10X-2P, Thorlabs, USA).

On the Bruker system, 500 × 500 μm (256 × 256 pixels) images were acquired at 3.6 Hz using a linear galvanometer raster scanning (Cambridge Research). Emitted photons were reflected by a 660 nm longpass primary dichroic mirror (660dcxxr, Chroma) and filtered by an infrared blocker (ET680sp-2P, Chroma). Green fluorescence was separated by the combination of a longpass dichroic beam splitter (T565lpxr, Chroma) and a bandpass filter (ET525/70m-2P, Chroma), and detected by a GaAsP photomultiplier tube (H7422PA-40 SEL, Hamamatsu Photonics). Images were digitized and recorded using PrairieView (Bruker, USA). On the Bergamo system, 500 × 500 μm (512 × 512) pixels images were acquired at 30 Hz using a galvo-resonant scanner (LSK-GR08, Thorlabs, USA). Emitted photons were filtered by a 750nm shortpass primary dichroic mirror (FRM2000, Thorlabs, USA). Green fluorescence was separated by a bandpass 525nm fluorescence filter (MF525-39, Thorlabs, USA), and detected by a GaAsP photomultiplier tube (PMT2100, Thorlabs, USA). Images were digitized and recorded using ScanImage (Vidrio Technologies, USA) software. In both systems, the beam power was controlled via a Pockels’ cell (Conoptics, USA) and was measured both at the laser output and after the objective lens to verify the power level at the brain.

Ultrasonic stimulation was delivered with the custom assembled ring transducer (allowing to image samples through its aperture), affixed below the objective via a 3D printed holder specifically designed to ensure the coaxial alignment of optical and acoustic foci. The same electrical drive chain (waveform generator and voltage amplifier) employed during calibration was used to drive the transducer, triggered by the acquisition software.

During experiments, mice were head-restrained on a custom-made holder and placed below the objective/transducer complex, affixed atop a dual axis goniometer stage allowing compensation for minor tilts in the cranial window. Transparent ultrasound gel (Aquasonic CLEAR, Parker Laboratories, USA) was deposited between the lens and the transducer, and between the transducer and the cranial window, to ensure optical and acoustic coupling. XYZ motors were then used to bring the objective and transducer above the cranial window, and to find regions of interest across the visual cortex and cortical depths.

During imaging acquisitions, ultrasonic perturbations were delivered as 200 ms long pulsed stimuli with a carrier frequency of 2.1 MHz and a constant pulse repetition frequency of 100 Hz. Multiple stimulus conditions were presented with various peak pressure amplitudes (from 0 to 0.8 MPa) and stimulus duty cycles (from 5 to 80%). For each condition, mice were presented with 16 consecutive stimuli with an inter-trial interval between 25 and 28 s. No more than 15 conditions were explored per session, resulting in a total imaging time below 2 hours.

### Analysis of calcium fluorescence movies

Acquired fluorescence movies were analyzed using custom-written Python code. High sampling rate (30 Hz) acquisitions were resampled to 5 Hz prior to processing and analysis. Initial movie inspection revealed the presence of two types of image-wide artifacts: an acquisition onset-locked fluorescence transient caused by the Pockels cell initialization, and a single frame, stimulus-locked image blurring caused by sonication. Each artifact resulted in isolated corrupted frames, which were replaced by either their preceding instance (for sonication frames) or by their following instance (for initial frames).

In the PV dataset specifically, signals were also contaminated by slow (<< 1 Hz) and large-amplitude (> 20%) variations in fluorescence occurring over the entire field of view; a physiological artifact that was likely caused by the hemodynamics of nearby vessels dynamically altering light absorption in the excitation/emission paths^49^. To remove this artefact, a global normalization strategy was employed. A reference frame (*F*_*ref*_) was identified for each movie using median projection, and each frame was then regressed against this reference to establish a global linear mapping (*F* ≈ *a* · *F*_*ref*_ + *b*). Subsequently, an inverse mapping (*F*^*^ = (*F* − *b*)/*a*) was applied to correct for global variations, effectively removing whole-frame fluctuations while preserving stimulus-evoked responses in identified neurons. Notably, this correction step was exclusive to the PV dataset, as the Thy1 and SST datasets did not exhibit similar signal contamination, likely due to their higher signal-to-noise ratios.

All pre-processed movies were denoised with a recursive prediction/correction algorithm based on the Kalman filter (https://imagej.net/ij/plugins/kalman.html) with a gain set to 0.5. The suite2p analysis pipeline^50^ with appropriate sampling rate (3.6 or 5 Hz) and sensor timescale (1.25 s for GCaMP6s, 0.7 s for GCaMP7f) settings was then used to align frames to a common session reference (via non-rigid registration), automatically identify neuronal somata within the field of view, and extract spatially averaged fluorescence time courses for each soma and its peri-somatic neuropil. Segmentation results were inspected by overlaying identified somata atop of the session reference image, and datasets displaying defective segmentation patterns were excluded. In parallel, registration metrics were used to quantify rigid motion within the imaging plane, and trials with significant stimulus-induced motion artifact were rejected.

Raw fluorescence traces were neuropil-corrected (*F*_*neuropil corrected*_ = *F*_*soma*_ − *α* · *F*_*neuropil*_) to remove background fluorescence. The scaling factor *α* was set to 0.7 (as in previous work^25^), or to the maximal value producing a non-negative output (so as to prevent over-correction). For each acquisition, the time-varying baseline *F*_0_ of each neuropil-corrected fluorescence signal was computed using a 30 s sliding percentile window. An adaptive percentile value was chosen for each neuron based on the overall signal skewness (as in previous work^51^), to compensate for differences in overall activity between neurons. For each imaged region, the stability of fluorescence intensity was inspected by computing the relative variation in the population-average fluorescence baseline across the session, and datasets with deviations from the mean larger than 50% were excluded from the analysis. Relative changes in fluorescence were computed for every neuron as *ΔF*/*F* = (*F*_*neuropil corrected*_ − *F*_0_)/*F*_0_. Finally, gaussians were fitted to the histogram distribution of ΔF/F traces to estimate the mean and standard deviation of the noise in these signals (*μ*_*n*_ and *σ*_*n*_, respectively) and compute z-scored traces as *Z* = (Δ*F*/*F* − *μ*_*n*_)/σ_*n*_.

Z-scored ΔF/F traces were then median-aggregated across valid trials for each cell and condition, and multiplied by the square root of the number of trials to retain z-score interpretability. For each cortical cell type, a global average of trial-aggregated z-scored traces was computed across all neurons, datasets, and stimulation conditions. The full width half maximum of the response waveform was then used to estimate the cell type’s “nominal” response duration.

For each neuron and sonication trial, pre- and post-stimulus activity levels were respectively calculated by averaging traces in a 1.4 s baseline window immediately preceding the stimulus and the cell-type-specific response interval. Response amplitudes were computed as the difference between these two averages. Response amplitudes larger than 1.95 (corresponding to a 5% significance level for unidirectional detection) were classified as TUS-evoked responses. Finally, cells showing statistically detectable responses across at least 50% of substantial exposure conditions (i.e., *I*_*SP,TA*_ ≥ 2 *W*/*cm*^2^) were classified as “TUS-responding neurons”.

For each imaged region, time traces and response amplitudes were aggregated across all identified cells for each condition to obtain descriptive statistics (mean and standard error), which were then aggregated across all datasets of each mouse line using a weighted average based on the number of cells identified in each dataset.

### Analysis of TUS-evoked response scaling laws

To evaluate the (mis-)alignment between pressure and DC sweeps for a given acoustic dose *γ* = (*P, DC*), we first projected parameter values from the pressure and DC sweeps onto the *γ* space to obtain projected response curves (*y*(*γ*_*P*_) and *y*(*γ*_*DC*_), respectively). We then fitted a sigmoid to the projected DC response curve (chosen because of its wider acoustic dose span) to obtain a predictive response model ŷ (*γ*), which we sampled at values covered by the pressure sweep. We then computed absolute differences between observed and predicted values and divided them by the standard errors of the observations to obtain a vector of normalized errors:

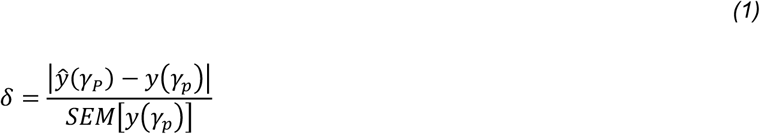

Error statistics (mean and standard error) were then computed for each cell type and dose projection.

### Computational model of TUS-evoked circuit responses in the visual cortex

To simulate and predict TUS-evoked responses to TUS stimuli, we developed a three-population rate model of local interactions between excitatory, SST and PV populations. In this model, the time-varying activity of each cortical population is captured by a system of first-order ordinary differential equations (ODE):

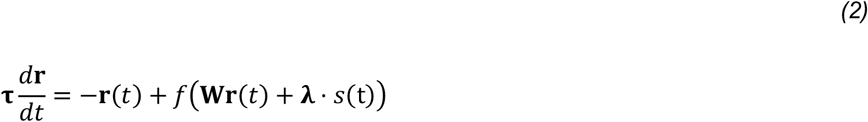

where:

- **r** is the vector of time-varying rates,
- **τ** is the vector of cell-type-specific decay time constants,
- *f* is an input-output activation function, defined as a rectified linear unit with specific threshold (*θ*) and slope (*g*) parameters,
- **W** is the population connectivity matrix of the model,
- **λ** is a vector of cell-type-specific stimulus sensitivities,
- *s* is a time-varying vector representing the ultrasound stimulus.

Cell-type-specific electrophysiological properties were extracted from the *Cell Feature Dataset* of the Allen Brain Atlas^27^, which was collected in acute slice preparations. We restricted our analysis to cells from the mouse primary visual cortex, layer 2/3. PV (n = 38) and SST (n = 16) cells were identified from mouse lines with corresponding Cre-tags (Pvalb-IRES-Cre and Sst-IRES-Cre, respectively), while excitatory cells (n = 38) were extracted from mouse lines containing exclusively cells with spiny dendritic structures (*Cux2-CreERT2, Esr2-IRES2-Cre, Glt25d2-Cre_NF107, Slc17a6-IRES-Cre, Scnn1a-Tg3-Cre, Scnn1a-Tg2-Cre*, and *Rbp4-Cre_KL100*). We averaged features from all constituent cells of each category to obtain a fixed set of time constants and activation parameters.

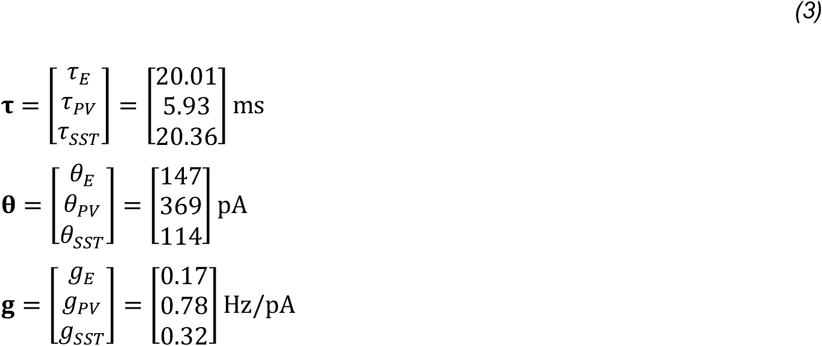

The reference network connectivity matrix was derived from the *Synaptic Physiology* dataset of the Allen Brain Atlas^31^, which was collected in acute slice preparations from the mouse primary visual cortex. We restricted our analysis to neurons from cortical layer 2/3. PV (n = 204) and SST (n = 251) cells were identified from mouse lines with corresponding Cre-tags, while excitatory cells (n = 249) were identified according to a morphological criterion (spiny dendritic structure). Synaptic connections identified from multi-patch recording recordings across these cells were used to derive a matrix of connection probability between each cell class. From the same dataset, we extracted the average post-synaptic potential (PSP) amplitude evoked in each cell class from both inhibitory and excitatory synapses, and broadcasted them according to pre-synaptic cell type to obtain a matrix of unitary synaptic strength between each cell class. Finally, following a similar approach as in previous work^28^, we multiplied the connection probability and unitary synaptic strength matrices to obtain our final set of network coupling weights.

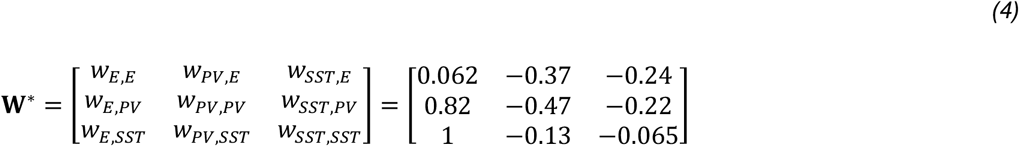

To simplify the model units, activation parameters were normalized to those of the excitatory unit used in a previous study^42^ (*θ*_*E*_ = 5, *g*_*E*_ = 1), while coupling weights were normalized to the E→SST value used in that same study (*w*_*E,SST*_ = 12).

The ultrasound stimulus (*t*) was defined as a 200 ms long square pulse with an amplitude equal to the stimulus I_SPTA_, in W/cm^2^, transformed into a model input via multiplication with the sensitivity parameter **λ**.

The model was implemented in custom-written Python code. Numerical integration was performed using a variable time step ODE solver with initial conditions set to **r** = 0. Dose-dependent response profiles were then extracted by simulating the model for a linear set of acoustic intensities spanning our experimental range (0 − 15.8 W/cm^2^) and extracting the steady-state activity reached by each population during the stimulus period.

We evaluated the accuracy of our model by comparing predicted response profiles (*y*^*^) against corresponding empirical references (*y*), which were obtained by sampling our cell-type-specific predictor functions at identical I_SPTA_ values. Since calcium fluorescence does not guarantee a consistent linear mapping to the underlying spiking activity across cells, animals, and mouse lines^25^, we chose to perform our comparisons in a normalized space. For each set of predicted response vectors, we quantified the model prediction error as:

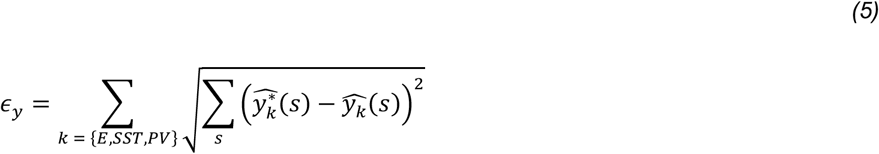

Where:

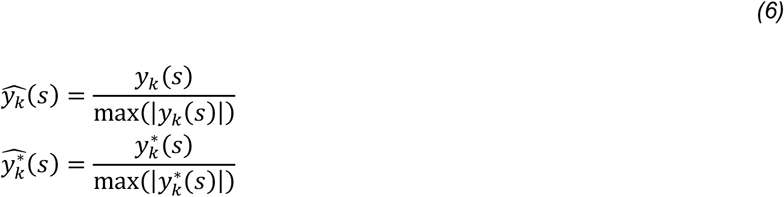

To find the set of network coupling weights and stimulus sensitivities for which the model best approached empirical data, we carried out a multi-dimensional parametric optimization with a differential evolution algorithm (*scipy*.*optimize*.*differential_evolution* function). The search bounds were set to [0, +20] for excitatory connections, [-20, 0] for inhibitory connections and [0, 5] for stimulus sensitivities. Since we did not strictly enforce inhibition dominance, the optimization algorithm inevitably explored parametric regions where stimulus-evoked responses did not reach stable steady-states, or where activity levels did not go back to baseline after the stimulus offset. These occurrences were respectively identified by thresholding linear regression slopes or average values on the activity vectors in appropriate time intervals, and sanctioned with an infinite penalty. Relatedly, some configurations produced response profiles that, despite providing reasonable agreement in the normalized space, differed by several orders of magnitude across populations. To avoid algorithmic convergence towards these configurations, we implemented a “disparity” penalty proportional to the maximal ratio of maximum activities reached across populations:

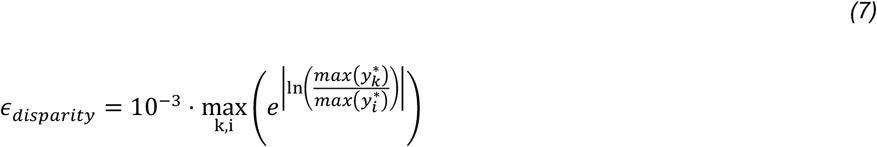

Note that the (logarithm → absolute value → exponential) transformation is used simply to ensure all ratios are ≥ 1. Finally, to avoid converging towards unrealistic network architectures, we enforced a penalty on absolute relative deviations of the tested matrix **W** from the reference connectivity matrix **W**^*^:

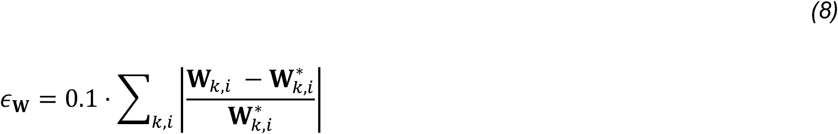

The resulting loss function for stationary solutions was:

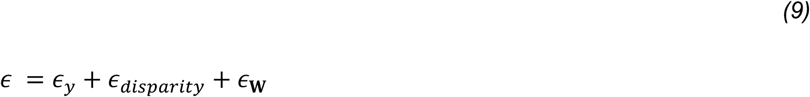

While differential evolution aims to find the global minimum of multivariate functions, it involves random initializations and stochastic mutations and could therefore converge towards different final solutions. Therefore, to evaluate the stability of our algorithmic convergence, we ran 5 parallel optimization runs for each model architecture and quantified the similarity between optimal solutions.

## ACKNOWLEDGMENTS

This work was funded by the NIH BRAIN Initiative (R01NS109885) and supported in part by an unrestricted grant from Research to Prevent Blindness to the NYU Department of Ophthalmology. T.L. was supported by a Postdoc.Mobility fellowship from the Swiss National Science Foundation (P500PB_211119). We thank Marco Petrozzi for experimental assistance, and Ben Stetler, Junhyook Lee, and Diego Asua for assistance with analyses. We are grateful to Dr. Wen-Biao Gan for providing the Thy1-GCaMP6s mouse line. We thank and Dr. Keith Murphy for helpful discussions.

## Author contributions

S.S., J.P.L. and R.C.F. designed the study and supervised the project. Y.Y. and J.P.L. built the experimental platform. T.L. mapped and calibrated the transducer. Y.Y., A.M.L. and J.P.L. performed surgeries. Y.Y., S.R.H.D., A.M.L. and C.G. acquired the experimental data. T.L. analyzed the data and performed model simulations. T.L., J.P.L. and S.S. wrote the manuscript. All authors discussed the results.

## Materials Availability

This study did not generate new unique reagents.

## Data and code availability

The pre-processed data and the code supporting the findings of this study will be made available upon publication.

## EXTENDED DATA

**Extended Data Figure 1.**
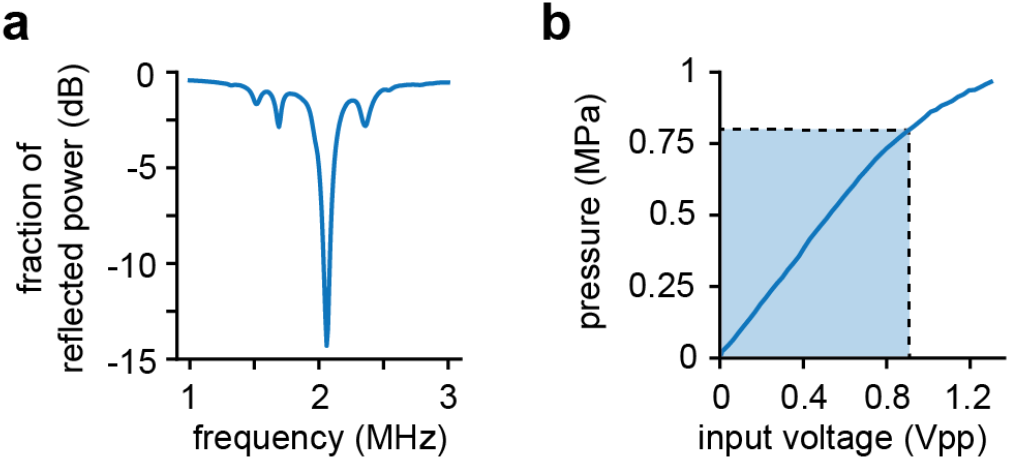
Transducer characteristics. **(a)** Fraction of input power reflected by the transducer as a function of driving frequency, measured by a network analyzer (E5061B, Keysight Technologies, USA). **(b)** Peak pressure amplitude measured at the transducer focus as a function of the input voltage of the signal generator (i.e., pre-amplification). The blue area indicates the range of input voltages (and corresponding pressure amplitudes) used during experiments.

**Extended Data Figure 2.**
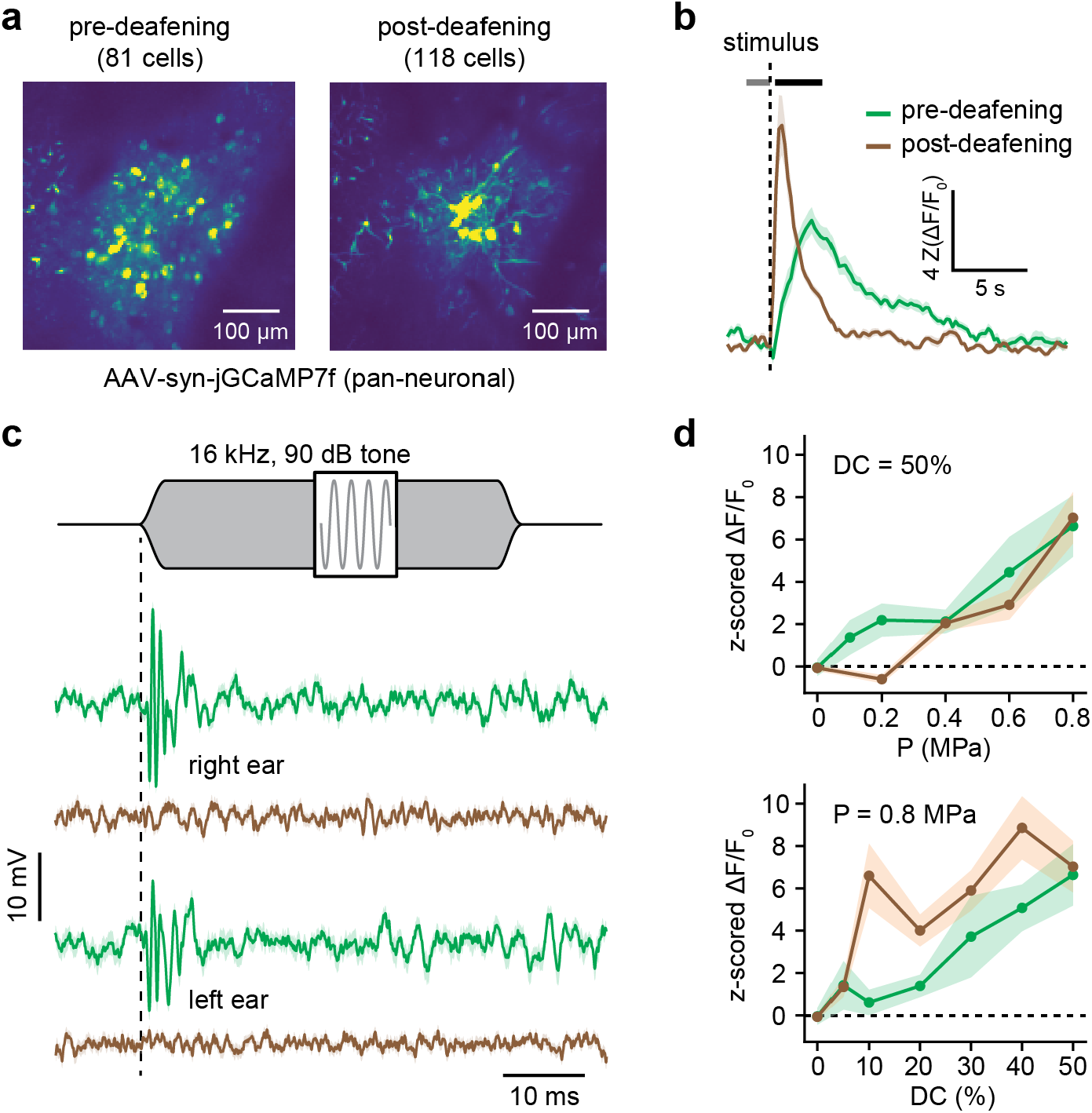
TUS-evoked responses in the auditory cortex before and after surgical deafening. **(a)** Maximal projection images (saturated at the 99^th^ percentile) of imaged fields of view in the mouse auditory cortex before (left, n=81 putative neurons) and after (right, n=118 putative neurons) surgical deafening. **(b)** Mean z-scored ΔF/F traces of TUS-evoked responses in identified TUS-responding neurons before (green, n=16 cells) and after (brown, n=21 cells) deafening, aggregated across stimulation conditions. Dark solid and light shaded areas indicate the mean +/-SEM of traces across selected neurons. Grey and black horizontal bars respectively indicate the pre- and post-stimulus windows used for the response quantification shown in (d). **(c)** Auditory brain stem responses (ABR) evoked by a 16 kHz, 90 dB tone directed to the right (top) or left (bottom) ear. Solid lines and shaded areas represented and mean +/-SEM of ABRs over 512 sweeps. **(d)** TUS-evoked response magnitude as a function of peak acoustic pressure (top, DC = 50%) and duty cycle (bottom, P = 0.8 MPa). Dark solid lines and light shaded areas denote the mean +/-SEM of response magnitudes across neurons.

**Extended Data Figure 3.**
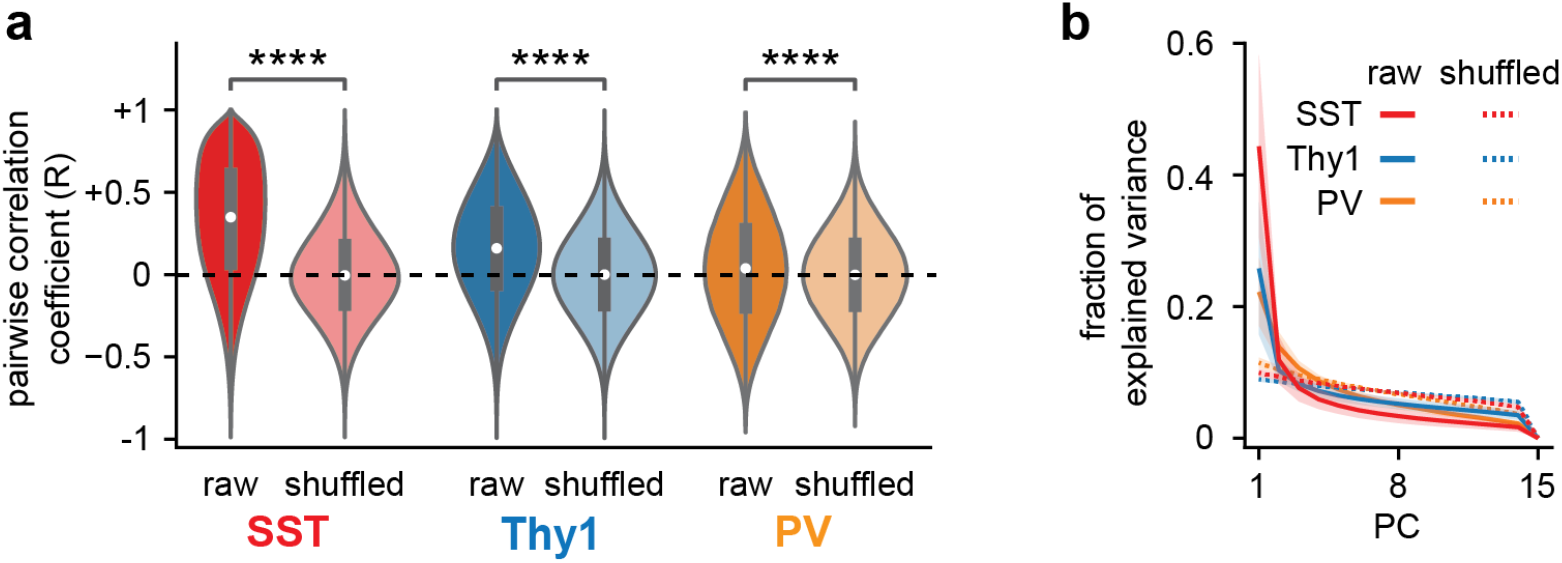
Cross-correlation and dimensionality analysis of TUS-evoked population responses. **(a)** Distribution of pairwise correlation coefficients of response vectors across cells, computed independently for each animal and stimulation condition. Stars indicate statistically significant differences between distributions computed from original and shuffled response vectors, for each neuron type (Mann-Whitney test, ****: p < 10^−4^). **(b)** Fraction of variance explained by each principal component following PCA decomposition of a 2D (cell, trial) population response matrix, computed independently across animals and acoustic doses. Solid and dashed lines denote the explained variance extracted from PCA on original and shuffled response matrices, respectively.

**Extended Data Figure 4.**
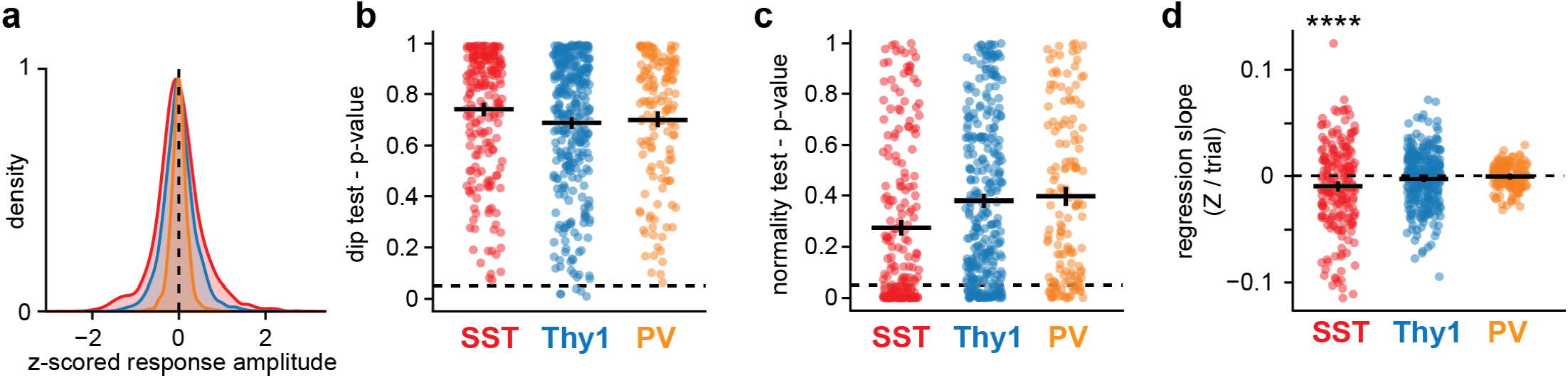
Trial-to-trial variability and stationarity of ultrasound evoked responses. **(a)** Density distribution of population average z-scored response vectors across animals and conditions, for each neuron type. **(b)** Distributions of p-values from Hartigan’s dip test for the unimodality of response vector distributions, for each neuron type, imaged region, and condition. Solid lines and error bars indicate the mean +/-SEM for each neuron type. The dashed line indicates the significance threshold for rejection of the null (unimodal distribution) hypothesis. **(c)** Distributions of p-values from the Shapiro-Wilk test for the normality of response vector distributions for each neuron type, imaged region, and condition. Solid lines and error bars indicate the mean +/-SEM for each neuron type. The dashed line indicates the significance threshold for rejection of the null (normal distribution) hypothesis. **(d)** Distribution of regression slopes fitted to z-scored vectors of population-average response magnitudes across trial sequences, for each imaged region and condition. Stars indicate distributions differing significantly from the null distribution (single sample t-test,==****: p < 10^−4^).

**Extended Data Figure 5.**
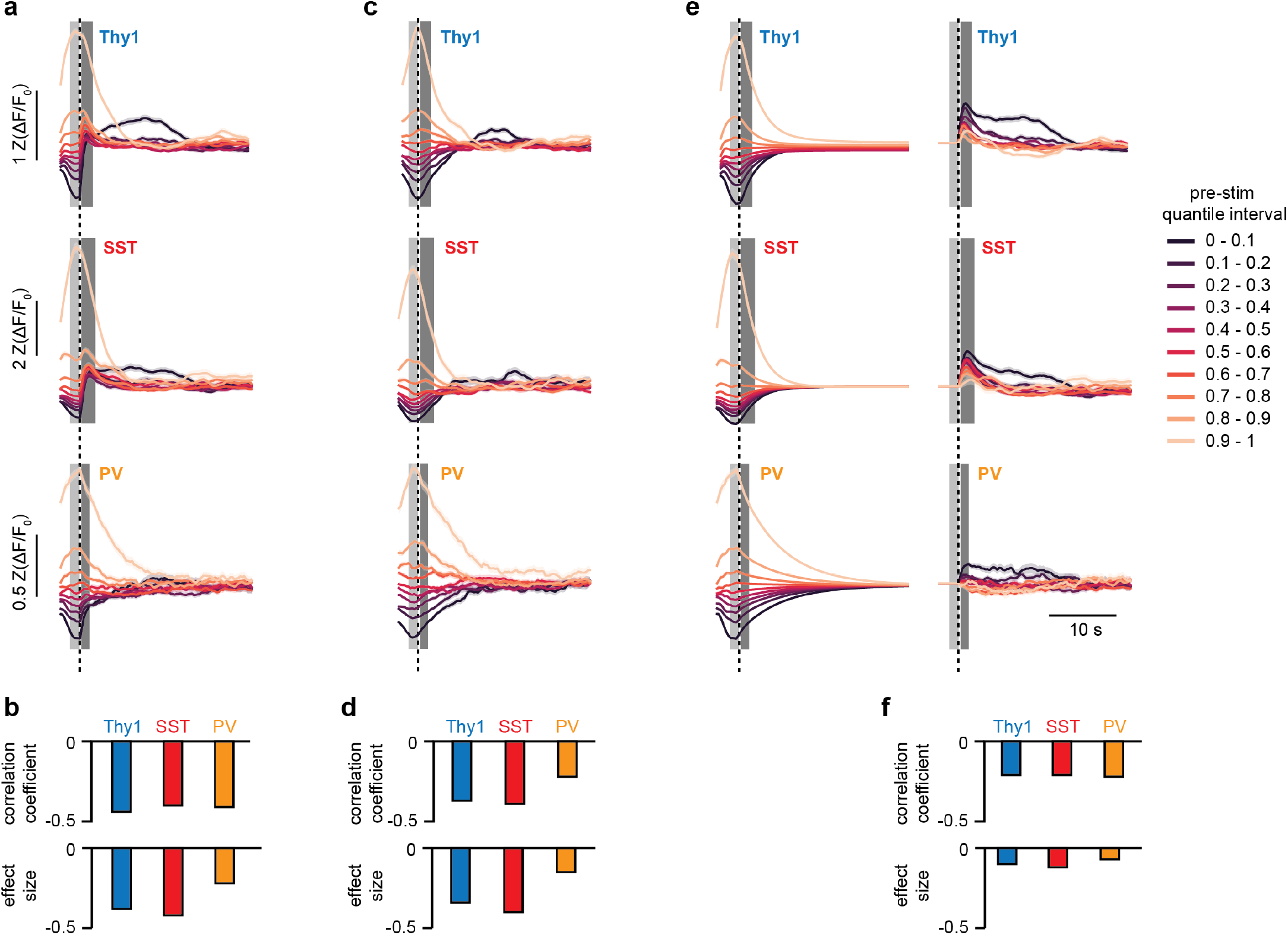
Dependence of TUS-evoked responses on ongoing neural activity. **(a)** TUS-evoked responses in Thy1 (top), SST (middle) and PV (bottom) populations, color-coded by levels of pre-stimulus activity. Population-average signals were binned into 10 separate groups based on their average level of activity in the pre-stimulus window. Traces and shaded areas represent the means +/-SEM in each quantile interval. The dashed line represents the time of stimulus, and the light and dark patches denote the pre-stimulus and post-stimulus analysis windows, respectively. **(b)** top: Spearman pairwise correlation coefficient between average pre-stimulus activity and average post-stimulus change in activity. Bottom: regression slope of average post-stimulus change in activity vs. average pre-stimulus activity. **(c)** Equivalent population-average traces from surrogate datasets generated by the iterative amplitude adjusted Fourier transform method. **(d)** Pairwise correlation coefficients (top) and regression slopes (bottom) between average pre-stimulus activity and average post-stimulus change in activity, computed from surrogate datasets. **(e)** Left: population-average traces where post-stimulus values are predicted from pre-stimulus values by an autoregressive (AR) model. Right: residual population-average traces obtained by subtracting the AR-predicted traces from the original traces. **(f)** Pairwise correlation coefficients (top) and regression slopes (bottom) between average pre-stimulus activity and AR-corrected average post-stimulus change in activity.

**Extended Data Figure 6.**
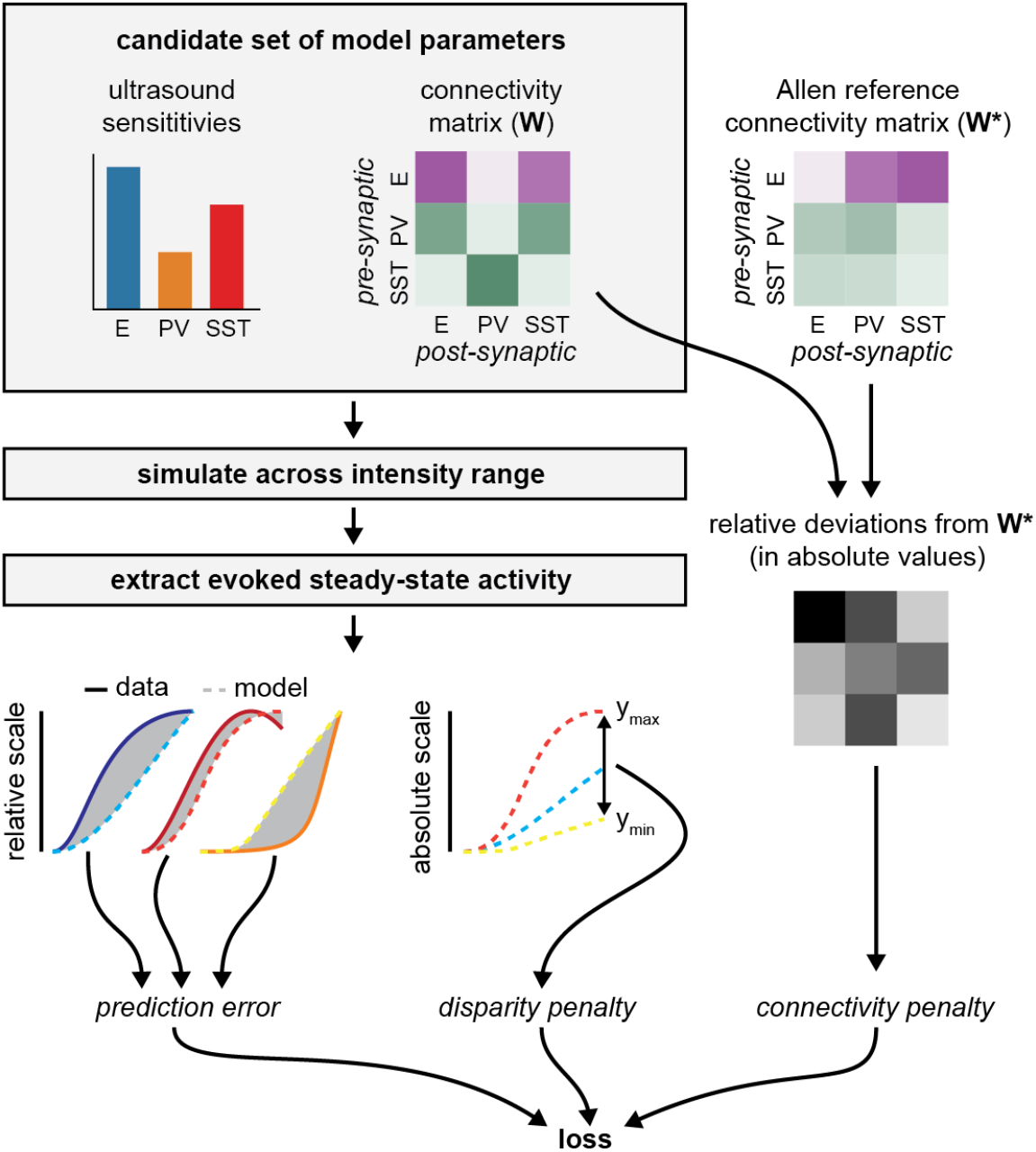
Schematic diagram of loss calculation for circuit model optimization. A candidate set of candidate parameters (intrinsic ultrasound sensitivities and synaptic coupling weights) is sampled. With this parameter set, the model is simulated across the empirical intensity range, and for each simulation evoked steady-state activities of each population are computed in the second half of the stimulus window (if applicable), to construct cell-type-specific response profiles. These profiles are then compared to empirical references on a normalized scale, yielding the prediction error. In parallel, the variation range of response amplitudes is computed for each population, and the maximal ratio of variation ranges across any 2 populations informs a disparity penalty. Finally, relative deviations of sampled coupling weights from the physiological reference from the Allen Brain Atlas are computed, and absolute values of these relative deviations are summed to inform a connectivity penalty. The final loss term is computed as a weighted sum of these 3 terms.

**Extended Data Figure 7.**
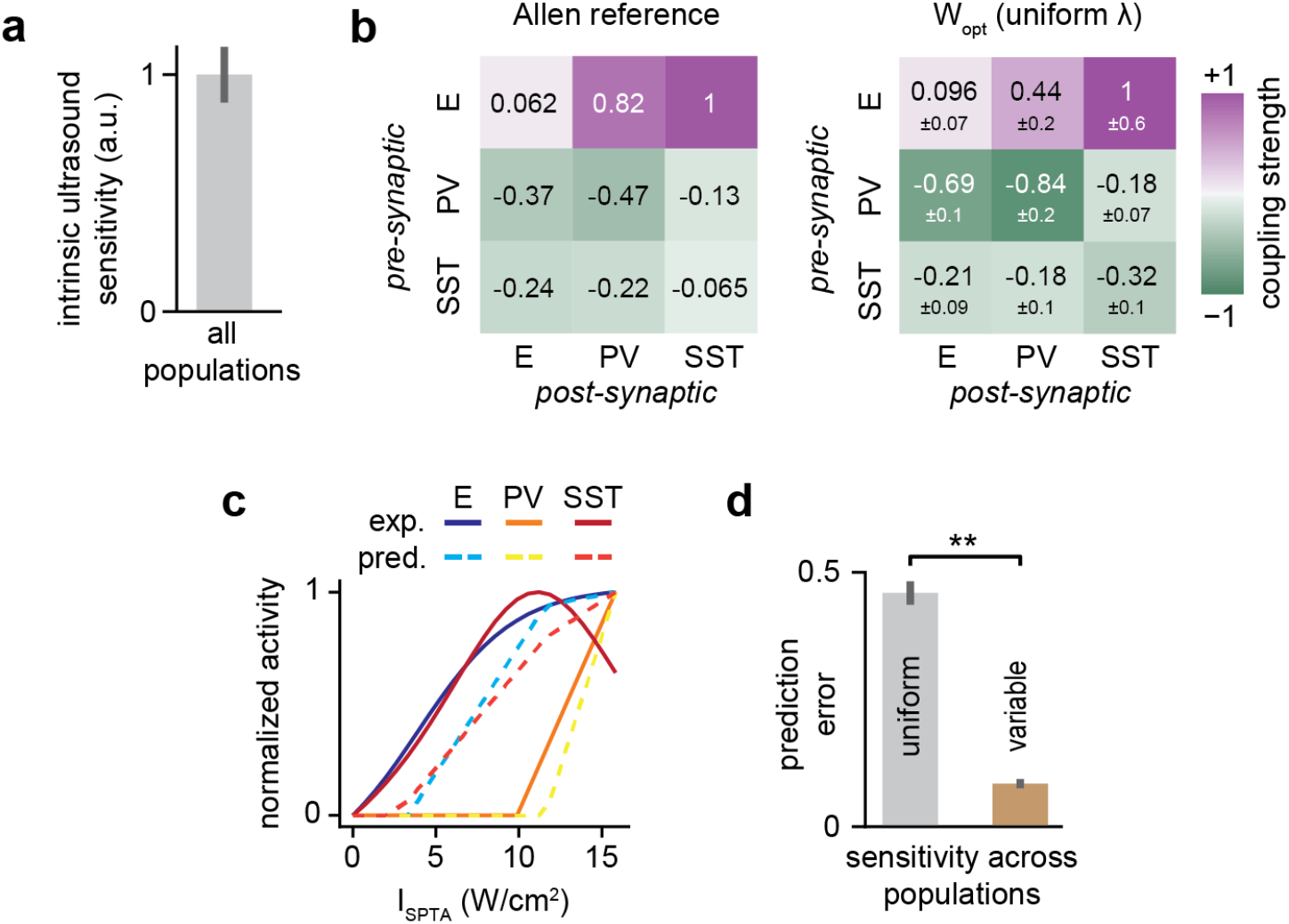
Impact of imposed uniform intrinsic ultrasound sensitivity across populations on model convergence and prediction performance. **(a)** Distribution of optimal stimulus sensitivities obtained across 5 model optimization runs. Bars and error bars depict the mean +/-SEM of sensitivities normalized to the unit interval. **(b)** Left: reference connectivity matrix of the mouse V1 micro-circuitry extracted from the *Synaptic Physiology* dataset of the Allen Brain Atlas. Excitatory (inhibitory) connections are depicted in purple (green), with a color saturation proportional to the coupling weight magnitude. Right: optimal connectivity matrix obtained across 5 model optimization runs. Mean +/-std values are indicated for each coupling weight. **(c)** Comparison of cell-type-specific and I_SPTA_-dependent response profiles predicted by the average optimized model against their empirical counterparts. Solid lines denote the empirically fitted predictors, while dashed lines denote the model predictions. Profiles were normalized to the unit interval. **(d)** Distribution of total model prediction errors (see Methods) for the I_SPTA_-dependent response profiles (left), compared to the standard case where constraints on uniform stimulus sensitivity are relaxed (right). Bars and error bars denote the mean +/-SEM of total prediction errors across 5 optimization runs. Stars denote statistically significant differences between distributions (Mann-Whitney test, **: p < 0.01).

**Extended Data Figure 8.**
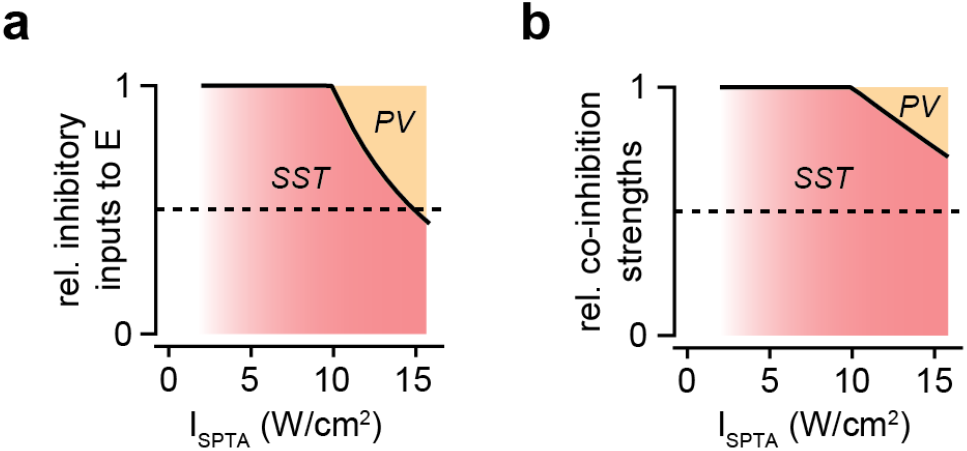
Dose-dependent SST and PV contributions to network inhibition predicted by the microcircuit model. **(a)** Proportion of inhibitory input to excitatory population coming from SST population, across the examined I_SPTA_ range. **(b)** Relative magnitude of SST->PV inhibitory input in the SST-PV mutual inhibition loop (i.e., SST->PV / (SST->PV + PV->SST)), across the examined I_SPTA_ range.

## SUPPLEMENTARY INFORMATION

### Derivation of acoustic dose functions

Prior equalities:

- the pulse duration (*PD*) comprises a finite number of acoustic periods (*f*^−1^):

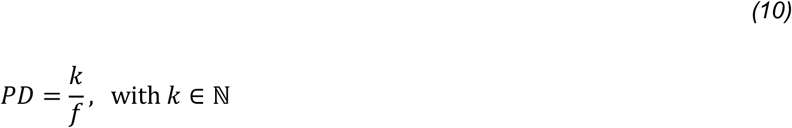
- the total burst duration (*BD*) comprises a finite number of pulse repetition intervals (*PRI*):

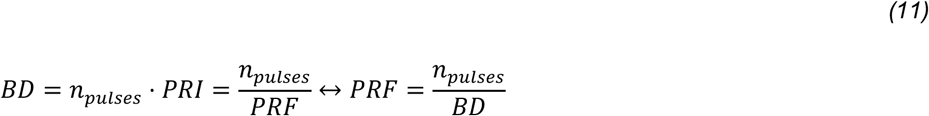
- Z = 1.6 MRayl is the medium’s specific acoustic impedance.

Dose functions derivation:

| Time-average<br>pressure amplitude | Time-average<br>acoustic intensity | Root mean square<br>acoustic intensity |
| --- | --- | --- |
| $P_{SPTA}$<br>$= \frac{1}{BD} \int_0^{BD} P_A(t) dt$<br>$= \frac{n_{pulses}}{BD} \int_0^{PD} P_A dt$<br>$= P_A \cdot PRF \int_0^{PD} dt$<br>$= P_A \cdot PRF \cdot PD$<br>$= P_A \cdot DC$ | $I_{SPTA}$<br>$= \frac{1}{BD} \int_0^{BD} I(t) dt$<br>$= \frac{1}{BD} \int_0^{BD} \frac{P^2(t)}{Z} dt$<br>$= \frac{n_{pulses}}{BD \cdot Z} \int_0^{PD} P^2(t) dt$<br>$= \frac{PRF}{Z} \int_0^{PD} (P_A \cdot \sin(2\pi ft))^2 dt$<br>$= P_A^2 \cdot \frac{PRF}{Z} \int_0^{PD} \sin^2(2\pi ft) dt$<br>$= P_A^2 \cdot \frac{PRF}{Z} \left[ \frac{t}{2} - \frac{\sin(4\pi ft)}{8\pi f} \right]_0^{PD}$ | $I_{SPRMS}$<br>$= \sqrt{\frac{1}{BD} \int_0^{BD} I^2(t) dt}$<br>$= \sqrt{\frac{1}{BD} \int_0^{BD} \left( \frac{P^2(t)}{Z} \right)^2 dt}$<br>$= \sqrt{\frac{n_{pulses}}{BD \cdot Z^2} \int_0^{PD} P^4(t) dt}$<br>$= \sqrt{\frac{PRF}{Z^2} \int_0^{PD} (P_A \cdot \sin(2\pi ft))^4 dt}$<br>$= \sqrt{P_A^4 \frac{PRF}{Z^2} \int_0^{PD} \sin^4(2\pi ft) dt}$<br>$= \sqrt{P_A^4 \frac{PRF}{Z^2} \left[ \frac{\sin(8\pi ft) - 8 \sin(4\pi ft) + 24 \pi ft}{64 \pi f} \right]_0^{PD}}$<br>$= \sqrt{P_A^4 \frac{PRF}{Z^2} \cdot \frac{24 \pi f \cdot PD}{64 \pi f}}$<br>$= \sqrt{\frac{3P_A^4}{8Z^2} \cdot PRF \cdot PD}$<br>$= \sqrt{\frac{3P_A^4}{8Z^2} \cdot DC}$<br>$= \frac{\sqrt{3}}{4Z} \cdot P_A^2 \cdot \sqrt{DC}$ |

### Predicted exposure metrics

For the maximal delivered acoustic dose (P_A_ = 0.8 MPa, DC = 80%):

- The mechanical index is computed as:

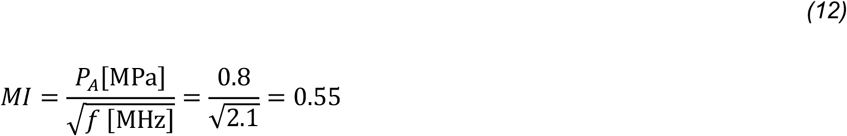

which is far below the regulatory limit of 1.9.
- The predicted temperature increase is calculated as:

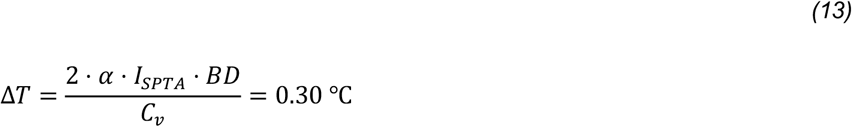

where C_v_ = 3.80 J/cm^3^/°C is the heat capacity per unit volume and a = 17.85 Np/m is the frequency-dependent acoustic absorption constant of brain tissue at 2.1 MHz^52^. This number is well within the physiological range of brain temperature variations.

